# SRS-FISH: High-Throughput Platform Linking Microbiome Function to Identity at the Single Cell Level

**DOI:** 10.1101/2021.07.23.453601

**Authors:** Xiaowei Ge, Fátima C. Pereira, Matthias Mitteregger, David Berry, Meng Zhang, Michael Wagner, Ji-Xin Cheng

## Abstract

One of the biggest challenges in microbiome research in environmental and medical samples is to better understand functional properties of microbial community members at a single cell level. Single cell isotope probing has become a key tool for this purpose, but the currently applied detection methods for measuring isotope incorporation into single cells do not allow high-throughput analyses. Here, we report on the development of an imaging-based approach termed stimulated Raman scattering - two-photon fluorescence *in situ* hybridization (SRS-FISH) for high-throughput structure-function analyses of microbial communities with single cell resolution. SRS-FISH has an imaging speed of 10 to 100 milliseconds per cell, which is two to three orders of magnitude faster than spontaneous Raman-FISH. Using this technique, we delineated metabolic responses of thirty thousand individual cells to various mucosal sugars in the human gut microbiome via incorporation of deuterium from heavy water as an activity marker. Application of SRS-FISH to investigate the utilization of host-derived nutrients by two major human gut microbiome taxa revealed that response to mucosal sugars tends to be dominated by Bacteroidales, with an unexpected finding that Clostridia can outperform Bacteroidales at foraging fucose.

## Main

With the rapid advances in both genotyping and phenotyping of single cells, bridging genotype and phenotype at the single cell level is becoming a new frontier of science^1^. Methods have been developed to shed light on the genotype-metabolism relationship of individual cells in a complex environment^2,3^, which is especially relevant for an in-depth understanding of complex microbial communities in the environment and host-associated microbiomes. For functional analyses of microbial communities, single cell isotope probing is often performed with nanoscale secondary ion mass spectrometry (NanoSIMS)^4–7^, microautoradiography (MAR)^8,9^, or spontaneous Raman microspectroscopy^10–12^ to visualize and quantify the incorporation of isotopes from labeled substrates. These methods can be combined with fluorescence *in situ* hybridization (FISH) using rRNA-targeted probes^13^, enabling a direct link between function and identity of the organisms. In addition, recently Raman-activated cell sorting has been developed, using either optical tweezers or cell ejection for downstream sequencing of the sorted cells^14–16^. While these approaches have expanded the possibilities for functional analyses of microbiome members^17^, all of the aforementioned methods suffer from extremely limited throughput. Consequently, only relatively few samples and cells per sample are analyzed via single cell stable isotope probing, hampering a comprehensive understanding of the function of microbes in their natural environment.

To overcome the limited throughput of Raman spectroscopy, coherent Raman scattering microscopy based on coherent anti-Stokes Raman scattering (CARS) or stimulated Raman scattering (SRS), has been developed^18,19^. Compared to CARS, the SRS signal is free of the non-resonant background and is linear to molecular concentration, thus permitting quantitative mapping of biomolecules^20,21^. Both CARS and SRS microscopy have successfully been applied for studying single cell metabolism in eukaryotes^22–25^. In a label-free manner, SRS imaging has led to the discovery of an aberrant cholesteryl ester storage in aggressive cancers^26,27^, lipid-rich protrusions in cancer cells under starvation^28^, fatty acid unsaturation in ovarian cancer stem cells^29^ and more recently in melanoma^30^. CARS and SRS have also been harnessed to explore lipid metabolism in live *Caenorhabditis elegans*^31–34^. Combined with stable isotope probing, SRS microscopy has allowed the tracing of glucose metabolism in eukaryotic cells^35,36^ and the visualization of metabolic dynamics in living animals^24^. Recently, SRS was successfully applied to infer antibiotic resistance patterns of bacterial pure cultures and D_2_O metabolism^24,37^. Yet, SRS microscopy has not yet been adapted for studying functional properties of members of microbial communities through the integration with FISH analyses.

Here, we present an integrative platform that exploits the advantages of SRS for single cell stable isotope probing together with two-photon FISH for the identification of cells in a high-throughput manner. To deal with the challenges in detecting a small number of metabolites inside small cells with diameters around a micron, we have developed a protocol that maximizes the isotope levels in cells and the SRS signal from the Raman band used for isotope detection. SRS is generated by near-infrared pulses while FISH is conventionally implemented using one-photon confocal fluorescence. To address this incompatibility, we developed a procedure for highly sensitive SRS metabolic imaging and two-photon FISH using the same laser source. These efforts collectively led to a high-throughput approach that enables imaging cell identity and metabolism at a speed of 10-100 milliseconds per cell. In comparison, it takes about 20 seconds to record a Raman spectrum from a single cell in a conventional spontaneous Raman-FISH experiment^38^.

In this work, we use the gut microbiome as a testbed for our technology. In the human body, microbes have been shown to modulate the host’s health^39,40^. Techniques looking into their activities and specific physiologies (i.e. phenotype) as a result of both genotype and the environment provide key information on how microbes function, interact with and shape their host. As a proof-of-principle, we used SRS-FISH to track the incorporation of deuterium (D) from D_2_O into a mixture of two distinct gut microbiota taxa. We show that SRS-FISH provides fast and sensitive information on the D-content of individual cells while simultaneously unveiling their phylogenetic identity. Subsequently, we applied this technique to complex microbial communities by tracking *in situ* the metabolic responses of two major human gut microbiome taxa to supplemented host-derived nutrients. Our study revealed that (i) Clostridia spp. could actually outperform Bacteroidales spp. at foraging on the mucosal sugar fucose and showed (ii) a significant inter-individual variability of responses of these major microbiome taxa towards mucosal sugars. Together, our results demonstrate the ability of SRS-FISH to unveil the function of particular microbes in complex communities, with throughputs that are two to three orders of magnitude higher than Raman-FISH and other metabolism-identity bridging tools, therefore providing a valuable multimodal platform to the field of single cell analysis.

## Results

### An SRS-FISH platform to link cell metabolism and cell identity

To retrieve information on the activity of single bacterial cells in culture or complex samples, we employed the D_2_O-based stable-isotope probing approach^10,37^. To this end, live cells present in simple (pure cultures) or complex (gut microbiome) samples were incubated in D_2_O containing-media, to enable incorporation of D into biomolecules of metabolically-active cells (**Fig. 1a**). Cells were subsequently fixed and subjected to FISH using fluorescently labeled oligonucleotide probes targeting rRNA, in order to reveal their phylogenetic identity (**Fig. 1a, b**). Samples prepared in this way were sequentially imaged to retrieve i) fluorescence signals from hybridized samples; and ii) chemical information that enables quantification of cellular D levels for the different taxa targeted by FISH (**Fig. 1b**). Measurements of D levels were performed using a dual output femtosecond (fs) laser that provided the pump (tunable from 680 nm to 1300 nm) and the Stokes (fixed at 1045 nm) beams. To quantify D incorporation, the pump beam was tuned to 852 nm to target the center of the C-D vibrational peak (2168 cm^−1^). The limit of SRS detection was determined to be lower than 4.3 mM for unlabeled dimethyl sulfoxide (DMSO) and 8.5 mM for D-labeled DMSO (DMSO-d6) by three standard deviations at zero concentration and the slope of the linear relationship between the SRS signal level and the concentration. (**Supplementary Fig. 1**). The SRS imaging resolution was determined to be around 300 nm in lateral and 2 µm in the axial direction, tested by 200 nm polymethyl methacrylate beads (**Supplementary Fig. 1**), and is thus suitable for measuring most bacteria. Taking into account the system sensitivity and SRS excitation volume, the minimum number of C-D and C-H bonds that can be detected by fs SRS are around five million and three million (**Supplementary Fig. 1**, see Methods). As bacterial cells have sizes that are comparable with the laser focus laterally and axially (**Supplementary Fig. 1**), the different volumes displayed by different bacterial species can influence the SRS intensity level. Therefore, in complex microbial communities, the C-D intensity of a cell alone cannot directly be translated into metabolic activity, especially when comparing between different groups of organisms. In order to normalize for the different volumes of cells in complex samples, the pump beam was tuned to 799 nm to target the center of the C-H bond vibrational peak (2946 cm^−1^) in the C-H stretch region. As the fs pulsed lasers have rather broad bandwidths, fs SRS has a total covering range of 200 cm^−1^ around the peak^28^. Thus, in this study, the C-D and C-H signature peaks at 2040-2300 cm^−1^ and 2800-3100 cm^−1^ can be mostly covered by fs SRS. We have observed that some bacterial species seemingly display signals at the silent regions (from 1800 to 2800 cm^−1^) that can be detected by SRS but not by spontaneous Raman spectroscopy (**Supplementary Fig. 2**). These signals may originate from processes other than SRS, such as transient absorption, photothermal lensing, and cross phase modulation^41,42^. To address this issue, off-resonance images were recorded with the pump beam tuned to 830 nm, targeting 2479 cm^−1^ in the silent region, and subtracted from both C-D and C-H SRS images (**Supplementary Fig. 2**).

**Figure 1.**
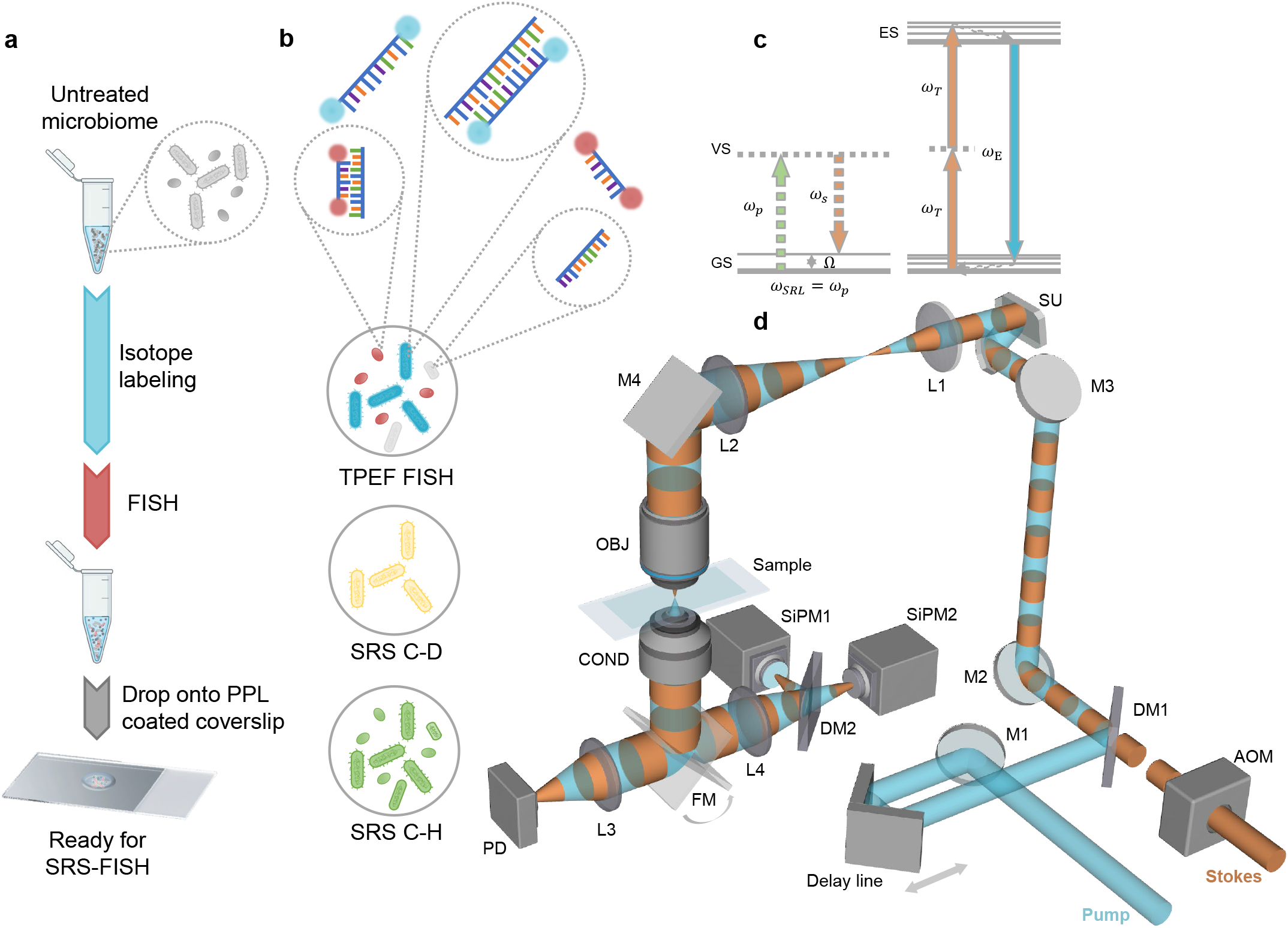
Stimulated Raman scattering (SRS)-fluorescence *in situ* hybridization (FISH) platform to bridge phylogenetic identity (genotype) and metabolic activity (phenotype) in microbes**. a**, Standard sample preparation process for SRS-FISH experiments. Pure bacterial cultures or complex microbiome samples are incubated in D_2_O-containing media to enable D incorporation into metabolically active cells. Samples are subsequently fixed and submitted to FISH. After FISH, samples are deposited in a glass cover slide and analyzed by SRS-FISH either directly while in the liquid environment, or after drying. **b**, Schematic representation of SRS-FISH imaging results. Samples are hybridized with fluorescently labeled oligonucleotide probes (double-labeled with either cyan or red fluorophores) targeting taxa of interest present in the sample. Fluorescence signal originating from hybridized samples (cyan and red) is then overlaid with SRS C-D signal (yellow) and SRS C-H signal (green) to reveal the metabolic activity levels of each identified bacterial cell. Organisms not targeted by the probes will not display fluorescence (grey). **c**, SRS and two photon excited fluorescence (TPEF) mechanism. ES: excited state. VS: virtual state. GS: ground state. *ω_P_*: pump beam laser frequency. *ω_S_*: Stokes beam laser frequency. *ω_SRL_*: stimulated Raman loss frequency. *Ω*: vibrational energy. *ω_T_* : TPEF excitation beam frequency. *ω_E_* : fluorescence emission frequency. **d**, SRS-FISH system. M1-M4: mirrors. AOM: acousto-optic modulator. DM1-2: dichroic mirrors. SU: scanning unit. L1-4: lenses. OBJ: objective. COND: condenser. FM: flip mirror. PD: photo diode. SiPM1-2: silicon photomultiplier.

FISH signals have been mostly detected by one photon microscopy, typically using widefield or confocal microscopy^43^. Since the near-infrared fs laser beams for SRS imaging fall into the excitation range of TPEF (**Fig. 1c**, right panel), we developed a two-photon FISH protocol. We focused on the detection of cyanine 3 (Cy3) and cyanine 5 (Cy5), two dyes that possess large two-photon cross sections^44^ and are commonly used in FISH studies due to their brightness and stability. Two silicon photomultipliers (SiPMs) mounted with 570 nm and 670 nm bandpass filters were used to selectively detect the fluorescence from Cy3 and Cy5, respectively (**Fig. 1d**). We confirmed that TPEF retrieves accurate fluorescence information with comparable imaging quality and speed as observed under a confocal microscope (discussed further below). Thus, two-photon FISH is a reliable tool for identity mapping, although with slightly lower resolution (∼300 nm) compared to confocal microscopy (usually ∼200 nm) (**Supplementary Fig. 1**).

To identify the targeted taxa and quantify their metabolic activity, the fluorescence signals from cells labeled by the hybridized probes are first acquired using TPEF (**Fig. 1b**). Then the same cells are imaged with SRS in both the C-D region that measures the extent of D incorporation and the C-H region, which allows normalizing for the influence of the cell volume on C-D intensity (**Fig. 1b**). Next, off-resonance SRS images were recorded and subtracted from both C-D and C-H images (**Supplementary Fig. 3**). Subsequently, single cell masks were created in CellProfiler^45^ using fluorescence images (or C-H images, in the case of pure cultures), and applied to all other channels, to enable precise quantification of D incorporation in FISH-targeted populations (**Supplementary Fig. 3**; detailed in Methods Section). Quantification of metabolic activity in single bacterial cells was represented by %CD_SRS_, calculated based on SRS intensities at C-D, C-H, and off-resonance, according to the formula: %CD_SRS_ =(I_CD_–I_off_)/(I_CD_+I_CH_–2 I_off_), where I stands for SRS intensity.

### SRS has sufficient sensitivity to track single metabolically-active bacteria tagged by FISH

Though SRS has been used for imaging isotope incorporation in bacterial and mammalian cells^24,37^, the impact of FISH on SRS measurements of hybridized bacterial cells has not been studied. For FISH, cells are fixed and subsequently hybridized at 46°C and washed at 48°C, and the buffers used in these steps contain formamide and detergent. These treatments have been previously shown to decrease both cellular material including lipids of the bacterial cells which can impact the D levels as assessed by spontaneous Raman^4,10,38^. To evaluate the impact of FISH on the SRS detection of cellular D levels, SRS signals from hybridized (FISH) and non-hybridized (no FISH) *Escherichia coli* cells grown in M9 and LB media containing various percentages of D_2_O and therefore displaying a wide range of cellular D contents were determined (**Fig. 2**). As expected, cells grown in complex LB media display lower levels of D incorporation compared to cells grown in M9 minimal medium containing equivalent percentages of heavy water (**Fig. 2**). The higher levels of D incorporation observed for growth on M9 (where only a defined carbon source is present) compared to a complex medium has been documented^46^. This is likely caused by the higher need for de novo biosynthesis of monomeric biomolecules, such as amino acids, nucleotides, or fatty acids, which are absent in minimal media but readily-available for direct uptake and incorporation from complex media. Despite these differences, for both media, we could observe a significant reduction of 19.72%±2.79% in the CD levels of hybridized cells (FISH) compared to fixed but non-hybridized cells (no FISH) (**Fig. 2b, e**, p<0.05, Mann-whitey U test). This reduction in %CD is smaller than reported for a similar comparison performed with spontaneous Raman measurements^10^ and might be explained by the fact that in the fixation and FISH protocols used here the contact time of cells with ethanol was reduced compared to the previously applied protocol (**Fig. 2c, f**).

**Figure 2.**
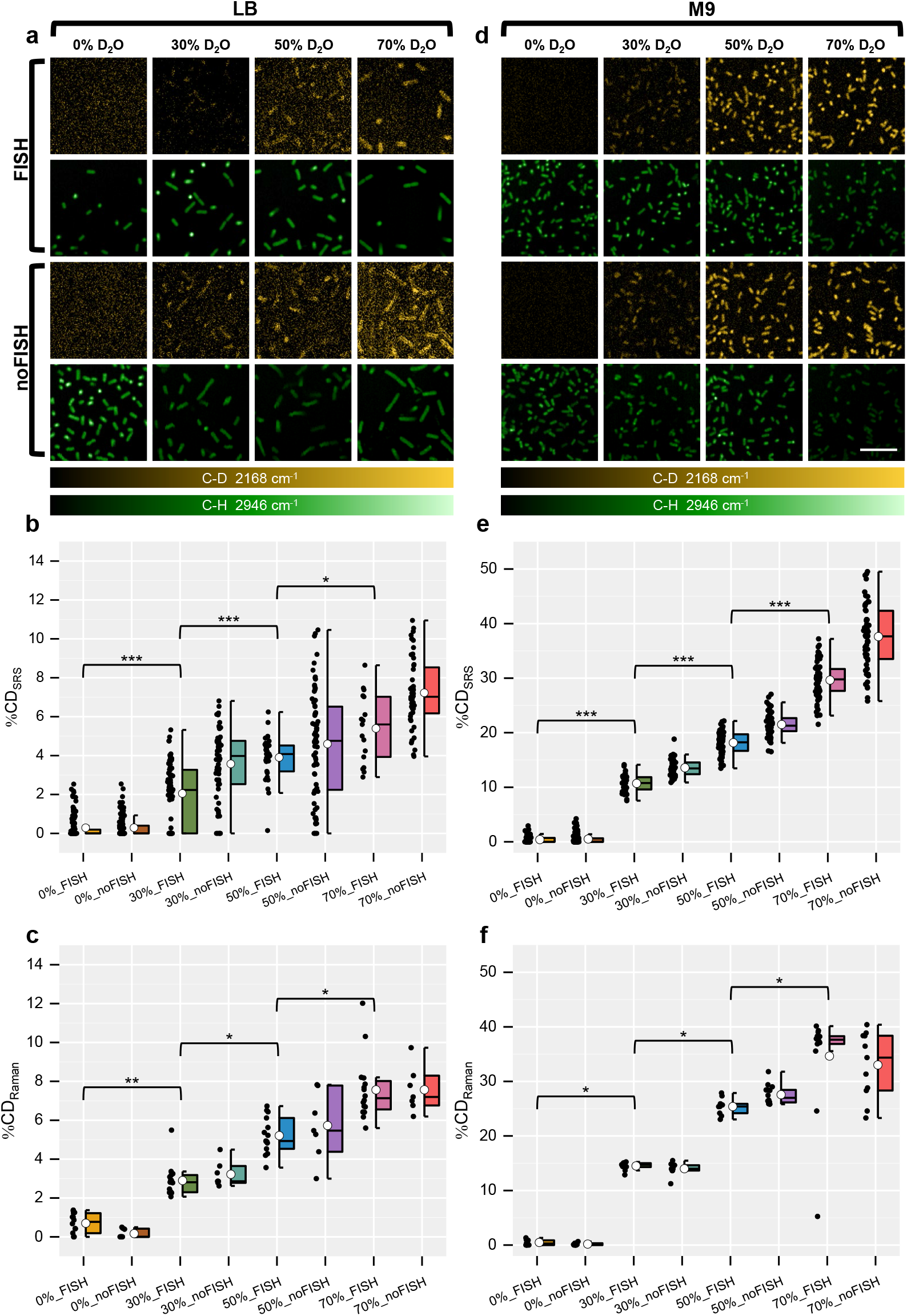
Sensitivity of the SRS-FISH platform to detect D_2_O metabolic incorporation into bacterial cells hybridized with oligonucleotide probes. SRS imaging of C-D (yellow) and C-H (green) intensities of single *E. coli* cells grown in LB (**a**) or M9 (**d**) media containing increasing percentages of D_2_O. For each condition, both hybridized (FISH, top panels) and non-hybridized (no FISH, bottom panels) cells were analyzed. Scale bar: 10 µm. Image contrast: (**a**) C-D channel, min 0 max 5 (arb. unit); C-H channel, min 0 max 30 (arb. unit). (**d**) C-D channel, min 0 max 1.5 (arb. unit); C-H channel, min 0 max 30 (arb. unit). Pixel dwell time: 100 µs. For details regarding data processing please refer to Supplementary Figure 3. **b, c**, Single cell C-D level distribution in *E. coli* cultures grown under the conditions described in **a**, measured with either SRS (**b**) or with spontaneous Raman microspectroscopy (**c**). NS: non-significant, p>0.05; *: 10^−5^<p<0.05; **: 10^−7^<p<10^−5^; ***: p<10^−7^ (two-sided Mann-whitney U test). **e, f**, Single cell C-D level distribution in *E. coli* cultures grown under the conditions described in **b**, measured with either SRS (**e**) or with Spontaneous Raman microspectroscopy (**f**). In **b, c, e, f**, each dot represents a cell. Boxes represent median, first and third quartile. Whiskers extend to the highest and lowest values that are within one and a half times the interquartile range. The white circle in the middle of the box represents the mean value of the data.

Notwithstanding the impact of FISH on calculated %CD_SRS_, SRS microscopy enabled efficient detection and discrimination of hybridized cells displaying a wide range of D levels, with mean %CD_SRS_ varying from as low as 2%, and up to 38% (**Fig. 2b, e**; p<0.05, two-sided Mann-Whitney U test). To validate the accuracy of fs SRS for bacterial activity quantification, we compared %CD_SRS_ with %CD measured by spontaneous Raman microspectroscopy (%CD_Raman_^10^). Under our conditions, SRS displayed similar sensitivity as spontaneous Raman (**Fig. 2c, f**).

However, SRS acquisition is two to three orders of magnitude faster than spontaneous Raman (10-100 ms per cell vs 20 s per cell)^38^. This high speed enables SRS measurements of a much large number of cells, therefore increasing the power of statistical analysis and throughput. In addition, we obtained similar results using an additional bacterial species (*Bacteroides thetaiotaomicron;* **Supplementary Fig. 4**), indicating that SRS-FISH is a fast and versatile platform to track metabolic activity in FISH-targeted bacteria.

### SRS is compatible with two-photon FISH to link function and identity

Cellular tagging by fluorescent dyes can lead to background signals detected by SRS, impacting the %CD_SRS_. This can occur when a fluorophore absorbs one photon from the pump and one photon from the Stokes beam, instead of two photons with the same energy. The simultaneous absorption of photons from two different beams causes a transfer of modulation from the Stokes to the pump, and interference with the SRS signal through a phenomenon known as non-degenerate two-photon absorption (ND-TPA). Although two-photon absorption can be calculated according to the absorption cross-section of recorded dyes, there is scant data of ND-TPA to estimate its influence on SRS. In our setup, we could not detect any interference on SRS attributable to the fluorophores in the sample, as SRS signals recorded from hybridized and non-hybridized *E. coli* cells grown in the absence of D_2_O were at the same level (**Fig. 2**). Consistently, we did not observe any significant differences in %CDs of hybridized compared with non-hybridized cells grown in the absence of D_2_O for the two different media tested (**Fig. 2b, e**; 0% D_2_O). Therefore, the lower %CDs obtained after FISH is likely due solely to loss of cellular D during the FISH process, as proposed before^10^, but not caused by interference of dyes used for FISH with SRS imaging.

We then evaluated the compatibility of using the 852 nm pump beam and the 1045 nm Stokes beam to excite the Cy3 and Cy5 dyes coupled to the FISH probes (**Fig. 3a**). The two SiPMs integrated into the system efficiently detected the emission of single D-labeled *E. coli* cells hybridized with a Gam42a-Cy5 oligonucleotide probe and of D-labeled *B. thetaiotaomicron* cells hybridized with a Bac303-Cy3 probe (**Fig. 3a, Supplementary Table 1**). For particular biological samples, it has been shown that the fs beams used for SRS may generate signals in the visible range, and therefore interfere with TPEF imaging^47^. However, only a negligible TPEF signal was detected from D-labeled bacterial cells that were not hybridized (**Fig. 3a**), and therefore we conclude that SRS does not significantly interfere with TPEF detection.

**Figure 3.**
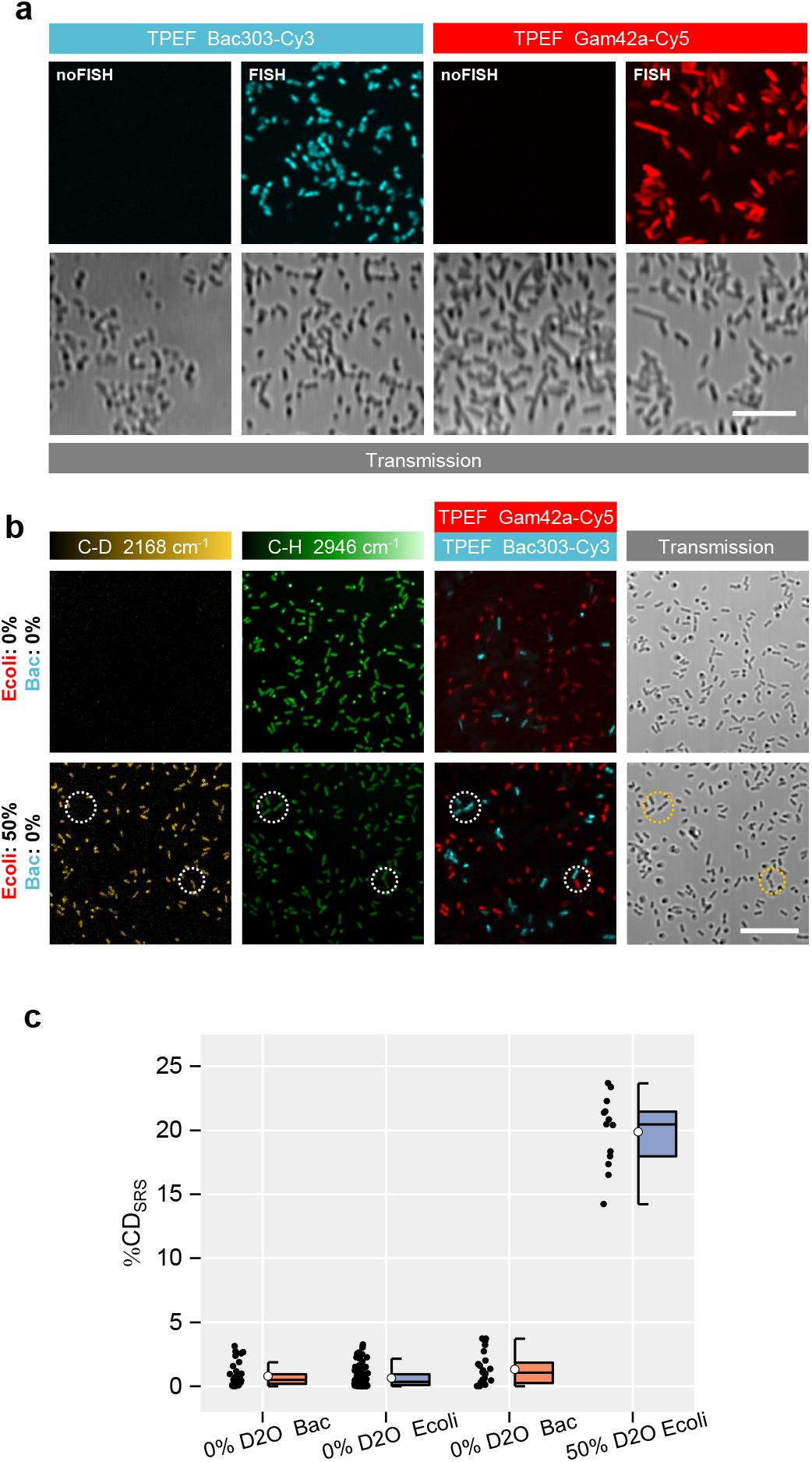
Genetic and chemical information can be acquired simultaneously by the SRS-FISH platform without interference. **a**, TPEF visualization of D-labeled *B. thetaiotaomicron* and *E. coli* cells hybridized (FISH) or not (no FISH) with Bac303-Cy3 (pseudo-coloured cyan) and Gam42a-Cy5 (pseudo-coloured red) oligonucleotide probes. No background in TPEF channels is detected for D-labeled, non-hybridized cells. Imaging under the dry conditions. Scale bar, 15 µm. Image contrast: Cy3: min 0.02, max 0.07 (arb. unit); Cy5: min 0.02, max 0.13 (arb. unit). **b**, Simultaneous SRS and TPEF imaging of artificial mixtures of unlabeled *B. thetaiotaomicron* and *E. coli* cells (top panel) or of D-labeled *E. coli* (50% D_2_O) and unlabeled *B. thetaiotaomicron* (0% D_2_O) cells (bottom panel). Bac: *B. thetaiotaomicron*. Ecoli: *E. coli.* SRS C-D (yellow) and C-H (green) intensities after off-resonance subtraction are shown for both mixtures. In both cases, cells were hybridized with the taxa-specific probes Bac303-Cy3 (cyan) and Gam42a-Cy5 (red), enabling cells from the two populations to be distinguished by TPEF imaging. Depicted circles highlight the presence of C-H, but absence of C-D signal in *B. thetaiotaomicron* cells identified by TPEF signal from the Bac303-Cy3 probe (in cyan). Scale bar, 15 µm. Image contrast: C-D channel, min 0 max 5 (arb. unit); C-H channel, min 0 max 30 (arb. unit). **c**, Single-cell C-D level distribution in the two different populations of the artificial mixtures presented in **b**, whose identity is revealed by TPEF imaging.

Having confirmed that SRS and two-photon FISH are compatible, we applied SRS-FISH to an artificial mixture of *E. coli* and *B. thetaiotaomicron* cells, each hybridized with taxa-specific oligonucleotide probes labeled with a different dye (**Fig. 3b**). In a first approach, both *E. coli* and *B. thetaiotaomicron* cells were grown in H_2_O-containing medium. Under these conditions, both cell populations display equally strong C-H signals, but no C-D signals were detected (**Fig. 3b**, upper panel). When we analyzed a mixture of D-labeled *E. coli* and unlabeled *B. thetaiotaomicron* cells, D signal only originated from *E. coli* cells targeted with the Gam42a-Cy5 probe, while *B. thetaiotaomicron* targeted by the Bac303-Cy3 probe displayed no C-D signal (**Fig. 3b**, bottom panel, highlighted circles). Using the signal obtained with TPEF and CellProfiler, we generated masks for either of the FISH-targeted populations (**Supplementary Fig. 3**), enabling automatic %CD_SRS_ calculation for each FISH-targeted population. While Cy5-tagged *E. coli* cells display %CDs of around 20, which reflects D incorporation from the D_2_O rich medium, Cy3-tagged *B. thetaiotaomicron* cells display %CDs close to 0, as expected (**Fig. 3c**). Collectively, these results demonstrate the feasibility of SRS-FISH to link metabolic active cells to their FISH-determined identity.

### High-throughput SRS-FISH for identifying mucosal sugar utilizers in the human gut microbiome

To demonstrate the applicability of the SRS-FISH approach to identify active taxa within a complex microbial community, we examined the responses of specific taxa from the human gut microbiota to additions of sugars from the mucus layer^48,49^ (**Fig. 4a**). Gut commensals able to forage on mucin can play a pivotal role in resistance to pathogen colonization and in modulating the host immune response^48,49^. In previous work, D_2_O combined with spontaneous Raman-activated cell sorting revealed that members of the families Muribaculaceae, Bacteroidaceae, and Lachnospiraceae are major mucosal sugar foragers in the mouse gut^17^. However, given the differences in microbiota composition of mice and humans, as well as differences in predominant types of mucus glycans that can be found in the two hosts^50,51^, it remains to be clarified if the same taxa are efficient mucosal sugar utilizers in the human gut, and what are their substrate preferences.

**Figure 4.**
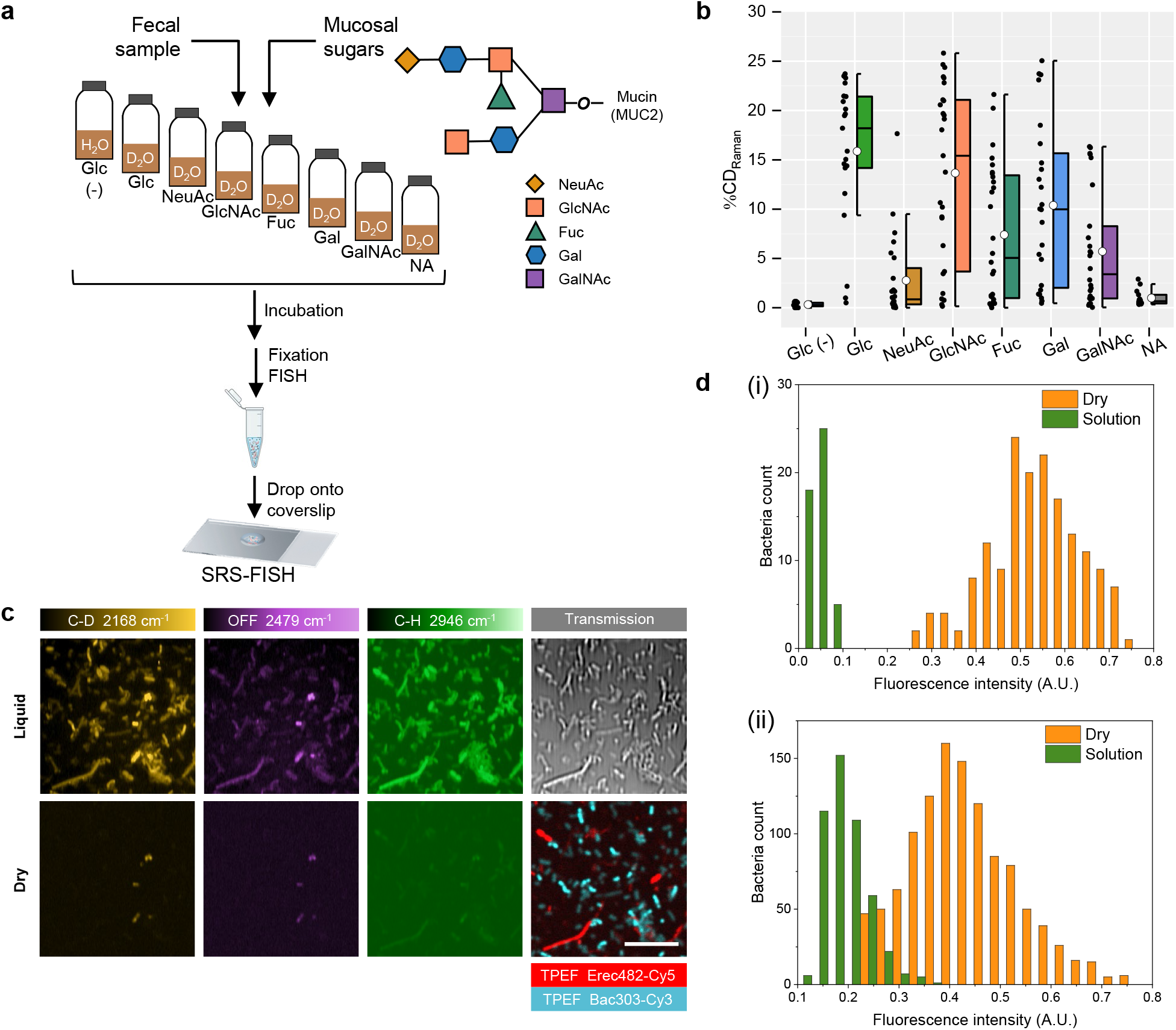
SRS-FISH to resolve the utilization of mucosal sugars by different microbiome taxa. **a**, Representation of the experimental setup. Freshly collected fecal samples were diluted in M9 medium and supplemented with different mucus *O*-glycan sugars in the presence of D_2_O. Negative- and positive-control incubations were performed in parallel by incubating the samples with Glucose (Glc) in the presence of H_2_O (labeled Glc (-)) or D_2_O (labeled Glc). Mucus *O*-glycan sugars include NeuAc: N-Acetylneuraminic acid, GlcNAc: N-Acetylglucosamine, Fuc: Fucose, Gal: Galactose, GalNAc: N-Acetylgalactosamine. A no amendment (NA) control with D_2_O but no sugar added was also included. Cells from all microcosms were processed as depicted (described in detail in the Methods section), and subsequently probed for D incorporation using spontaneous Raman, or D incorporation into taxa of interest using SRS-FISH. **b**, Single cell C-D level distribution of randomly selected microbiome bacteria supplemented with the different mucosal sugars and controls as described in **a**, measured by spontaneous Raman microspectroscopy. Each dot represents a cell. Boxes represent median, first and third quartile. The white circle in the middle of the box represents the mean value of the data. Whiskers extend to the highest and lowest values that are within one and a half times the interquartile range. **c**, SRS imaging of C-D (yellow), C-H (green) and off-resonance (purple) frequencies of hybridized gut microbiome cells when imaged in the liquid environment (top panel) or dry (bottom panel). TPEF signal from cells hybridized with the oligonucleotide probes Bac303-Cy3 and Erec-Cy5 is shown in cyan and red, respectively. Scale bar, 15 µm. For details regarding data processing please refer to Supplementary Figure 3. **d**, Intensity distribution histogram of fluorescence signals originating from cells hybridized with Cy3- (i) and Cy5-(ii) labeled probes, when imaged in solution or dried on a cover slide with confocal microscopy.

For this purpose, freshly collected human fecal samples were incubated with the five different mucin *O*-glycan sugars (N-acetylneuraminic acid: NeuAc, N-acetylglucosamine: GlcNAc, N-acetylgalactosamine: GalNAc, fucose, and galactose) in M9 minimal medium (without glucose) containing 50% of D_2_O (**Fig. 4a**). Under these conditions, we observed that human gut microbes responded to the mucosal sugars amended by incorporating D into their biomass, as well as to glucose (used as a positive control), as revealed using spontaneous Raman (**Fig. 4b**). Of note, only negligible incorporation of D was detected for cells that had been incubated in the presence of D_2_O but in the absence of any amended sugar, reflecting low metabolic activity driven e.g. by storage products or substrates released from decaying bacteria (**Fig. 4b**).

Having confirmed that human gut microbes were stimulated by mucosal sugars, we proceeded to identify which taxa respond to specific sugars using the SRS-FISH platform. For this purpose, oligonucleotide probes targeting two major groups of organisms in the human gut were applied: Bac303-Cy3 targeting *Bacteroides* and *Prevotella*^52^, among other Bacteroidales, and Erec482-Cy5 targeting members of the family Lachnospiraceae (also denominated *Clostridium* clusters XIVa and XIVb^53^) (**Supplementary Table 1**). These taxa were chosen because they constitute two of the most dominant and widespread phylogenetic groups of microbes in the human gut^54,55^. Additionally, a large percentage of organisms identified as efficient mucosal sugar foragers in the mouse gut are targeted or are closely related to organisms targeted by these probes^17^, and a large proportion of Bacteroidales spp. and Clostridia spp. have been shown to carry genes for mucosal sugar catabolism^56^. A major challenge encountered while imaging complex gut microbiome samples by SRS-FISH was that the TPEF signal from fluorescently labeled cells bleached much faster than what we previously observed in pure cultures. This might be explained by the relatively lower activity of many members of the gut microbiome, leading to a lower ribosome content and FISH signal when compared with the analyzed bacterial cultures, or by the lower cell wall permeability of particular microbiome members to oligonucleotide probes. To overcome this limitation, we acquired the TPEF signals from microbiome samples in a dried state, without any liquid between the coverslips that carried the sample (**Fig. 4c**, bottom panel). Stronger and more stable fluorescence signals than in liquid conditions were obtained for both Cy3 and Cy5-labeled cells (**Fig. 4d**). However, the SRS signal under dried conditions is weaker due to numerical aperture degradation caused by the refractive index mismatch (**Fig. 4c**, bottom panel, **Supplementary Fig. 1**). We have therefore optimized our protocols to first acquire the TPEF signal from the sample in a dried form, followed by the addition of water to the sample and SRS imaging of the sample in the liquid environment (**Fig. 4c**, upper panel). Importantly, the percentage of the fluorescently labeled cells targeted by each of the probes in the samples, determined either by TPEF (under the dry conditions) or by confocal microscopy, was in close agreement (18.7% versus 22.9% for Bac303-Cy3, respectively, and 77.1% versus 81.3% for Erec482-Cy5, respectively; **Supplementary Fig. 5**). In addition, we did not detect any cell-specific fluorescent signals from samples that had been hybridized with a mixture of Cy3- and Cy5-labeled negative control probes (**Supplementary Table 1, Supplementary Fig. 5**). These data suggest that the TPEF signal detected from Bac303-Cy3- and Erec482-Cy5-hybridized microbiome samples under dried conditions is accurate.

Using the SRS-FISH protocol optimized for complex microbiome samples, we examined the response of cells targeted by the Bac303-Cy3 and Erec482-Cy5 probes to each mucosal sugar in fecal samples from three different volunteers (**Fig. 5, Supplementary Fig. 6**). A first observation from our study was that the response to mucosal sugars differs from sample to sample (One-Way Anova test, p=7.863×10^−191^) (**Supplementary Fig. 7**), in both qualitative (discussed below), and quantitative terms. Quantitatively, we show that the overall microbiome response to the amended sugars (in terms of the number of active cells and their %CDs) was the highest for volunteer 2, while lowest cellular activity was detected for volunteer 3 (**Supplementary Figs. 7, 8**). This is not surprising, given that the human microbiome is highly individualized, and also that different fecal samples have been reported to contain different portions of viable cells^57,58^. Nevertheless, across the 3 samples the highest average number of active cells (and higher %CDs) was recorded in response to the mucosal sugar GlcNAc, followed by the response to galactose (**Supplementary Figs. 7, 8**), which is in agreement with the results obtained by spontaneous Raman (**Fig. 4b**). Overall, for all the samples analyzed we could detect a significant response of both targeted taxa to the mucosal sugars, with the exception of Erec482-targeted taxa to GalNAc in the sample from volunteer 1 (**Fig. 5 b-d, Supplementary Table 2**). In none of the samples did the no amendment control lead to the stimulation of a significant number of cells (**Fig. 5 b-d, Supplementary Table 2**). Furthermore, the response of Bac303-targeted taxa was overall higher than the response from Erec482-targeted taxa across all volunteers for all sugars tested, with the exception of fucose, where the inverse was observed for two of the volunteers (**Fig. 5**; **Supplementary Fig. 8**). These findings hold even after taking into consideration that the no specific signals in the C-D region were higher in the control group (H_2_O) for Bac303-targeted cells than for Erec482-targeted cells (**Fig. 5**).

**Figure 5.**
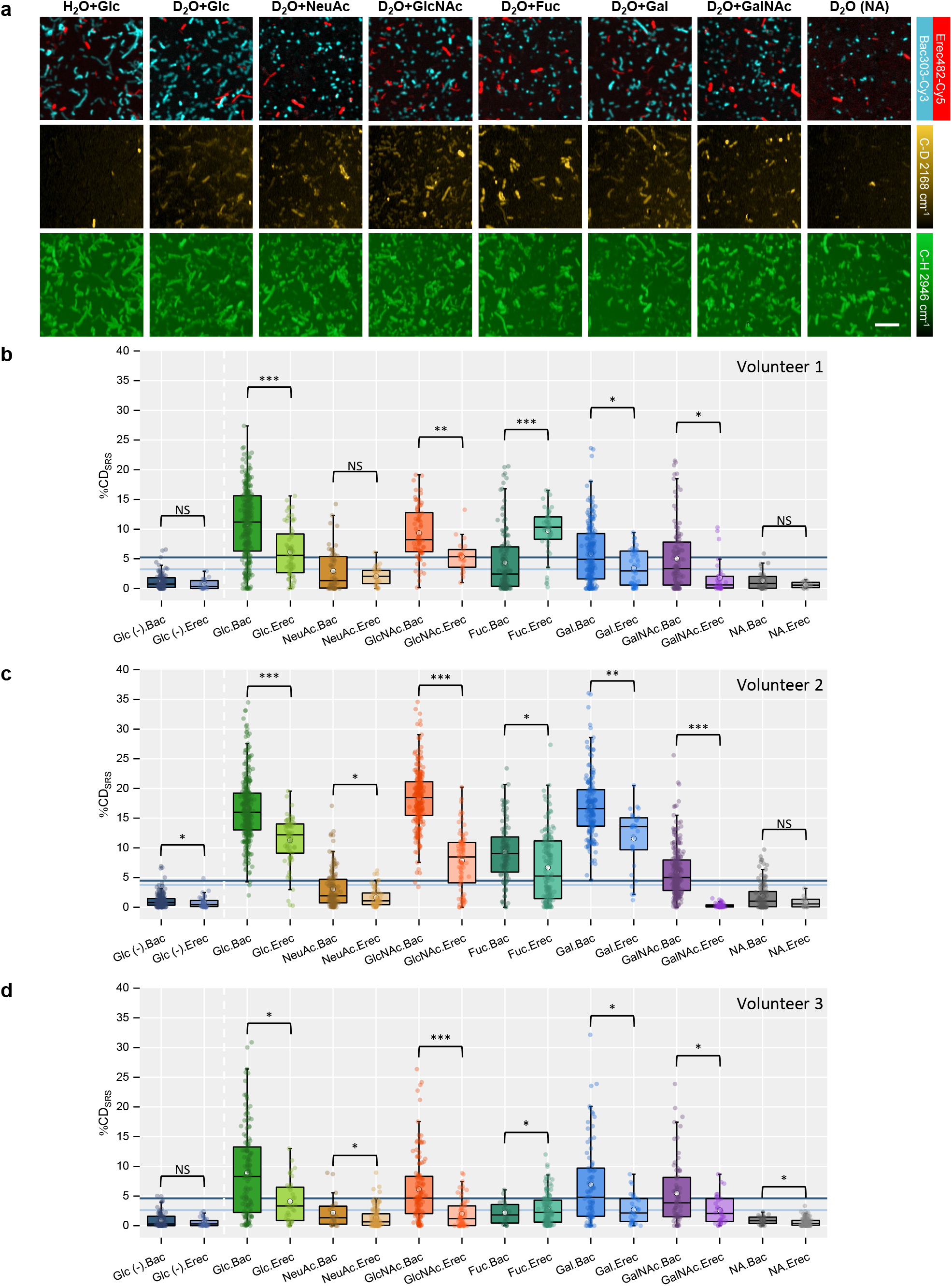
SRS-FISH uncovers the ability of major gut microbiome taxa to forage on different mucosal sugars. **a,** Microbiome samples of volunteer 1 were incubated with different mucosal sugars and hybridized with the oligonucleotide probes Bac303-Cy3 (cyan) and Erec482-Cy5 (red), respectively. Representative images obtained by TPEF (top row) and SRS (C-D middle row, C-H bottom row) are shown. Negative control: H_2_O+Glucose. Positive control: D_2_O+Glucose. NA: no amendment. Scale bar, 10 µm. For details regarding data processing please refer to Supplementary Figure 3. **b, c, d**, Single-cell C-D level distribution in the two different targeted taxa presented in **a**, measured by SRS for samples from 3 different volunteers. Box plots represent the median, first and third quartile with the extended lines represent the minimum and maximum value within 1.5 interquartile range from the first and third quartile. The white circle in the middle of the box represents the mean value of the data. The deeper grey blue line (mean +3SD of Bac303-Cy3 labeled cells in negative control) is the baseline over which Bac (Bac303-Cy3) cells are considered labeled. The lighter grey blue line (mean +3SD of Erec482-Cy5 labeled cells in negative control) is the baseline over which Erec (Erec482-Cy5) cells are considered labeled. The two-sided Mann-Whitney U-test was applied to compare the statistical significance between Bac and Erec bacteria for each amendment. NS: non-significant, p>0.05; *: 10^−5^<p<0.05; **: 10^−7^<p<10^−5^; ***: p<10^−7^ (two-sided Mann-Whitney U test).

## Discussion

Microbial communities are fundamental to the functioning of all ecosystems and the health of animals, plants, and humans. These microbiomes are typically investigated by meta-omic analyses that generate valuable annotation-based hypotheses regarding the metabolism of their members but are not suited for testing these hypotheses as gene annotations are often missing, wrong or incomplete^59^. Furthermore, many microbes have cell cycles, show considerable phenotypic diversity within isogenic strains, and the activity of microbes is influenced by their spatial arrangement in their habitat. Thus, there is an urgent need for direct functional analyses of microbes within complex samples with single cell resolution.

SRS-FISH fills a gap among the available tools linking function and identity in complex microbial communities due to its exceptionally high-throughput (10-100 millisecond per cell) and a sensitivity comparable to conventional spontaneous Raman-FISH. Overall, SRS-FISH is at least two orders of magnitude faster than state-of-the-art methods: MAR-FISH (2-20 days per sample)^60^, Raman-activated microbial cell sorting (>7.22 seconds per cell)^14^, FISH-nanoSIMS (>10 seconds per cell)^61^, which does not account for the long preconditioning time) and Raman-FISH (∼20 seconds per cell)^38^. Furthermore, implementation of FISH by TPEF can be advantageous when imaging thick biological specimens or live organisms, as near-infrared excitation enables deeper penetration into biological samples and causes less damage to cells^43^.

The application of SRS-FISH to the gut microbiome demonstrated the suitability of our approach to link identity to function within complex microbial communities, and at the same time revealed interesting new findings related to mucosal sugar foraging in the human gut. SRS-FISH measurements showed that Bacteroidales spp. tend to dominate the response to mucosal sugars over Clostridia spp. in all of the tested individuals (**Fig. 5, Supplementary Fig. 8**). Indeed, the notion that *Bacteroides* spp. are major mucus degraders has been demonstrated by several studies^17,56,62^. However, our results revealed that organisms from the *Clostridium* clusters XIVa/XIVb also substantially contribute to mucosal sugar degradation. Further, we show that larger fractions of Clostridia cells can forage on fucose compared to Bacteroidales cells (**Fig. 5, Supplementary Fig. 8**). Fucose is an important sugar in the colon as it occupies a terminal position on host glycans, thus being at the interface of microbiota–mucus interactions^63^. About 20% of human individuals naturally lack a functional copy of the *FUT2* gene, and thus lack almost all gut fucosylation^64^. Genome-wide association studies have shown that these individuals have increased susceptibility to inflammatory diseases linked to the gut microbiota, such as Crohn’s disease^65,66^. Additionally, mice that lack the Fut2 enzyme have simpler gut microbiomes that are accompanied by a decrease in unclassified Clostridiales^66^. These findings, together with our results, suggest that Clostridia may have been overlooked as fucose degraders in the gut^48^. Elucidating which particular Erec482-targeted organisms use fucose may be key to design individualized probiotic interventions aiming to restore the homeostasis in humans lacking *FUT2*, and therefore in reducing their predisposition to gastrointestinal disease.

Another interesting finding from our study is that the pattern of mucosal sugar foraging differs between the murine and human microbiome: human gut bacteria preferentially metabolize GlcNAc (**Fig. 5, Supplementary Fig. 7**), while the preferred sugar of the murine microbiome is galactose^17^. This could reflect the different overall compositions of human and murine colonic mucins, i.e., while the human colonic mucin carries predominantly GlcNAc-containing core 3- and core 4-based *O*-glycans, the murine colonic mucin is mostly characterized by galactose-containing core 1- and 2-type structures^50,51^. This finding may have important implications when translating results from mouse studies into humans.

There are several opportunities to further improve our SRS-FISH platform. These improvements include SRS-selective scanning of FISH targeted cells, which can even further improve the throughput of SRS-FISH when the taxa of interest appear in very low abundance. Besides, laser equipment with upgraded wavelength tuning speed will also provide the potential to gain higher throughput^67^. Other than the throughput, the sensitivity and resolution of SRS-FISH can also be improved by implementing visible SRS^68,69^. Of note, the excitation beam in visible SRS can efficiently excite fluorophores from cells targeted by FISH and help avoid the TPEF bleaching issue when imaging cells in a liquid environment. On the other hand, the number of taxa simultaneously tracked by SRS-FISH can be substantially increased using spectral unmixing and custom-designed FISH probes^70–72^. Regarding metabolism probing, besides using D_2_O as an activity marker to induce C-H peak shifts, deuterium-labeled substrates and other stable isotopes, such as ^13^C and ^15^N, could be used to track the metabolism of particular compounds and provide information on major catabolic activities and pathways. By targeting peaks between 400-1800 cm^−1^, SRS could fingerprint major intracellular macromolecules that display shifts due to the incorporation of stable isotopes. This could potentially be achieved by the implementation of hyperspectrum SRS with ultrafast delay-line tuning and machine learning into the SRS-FISH platform^16,73^. SRS-FISH would also be a useful tool to probe the distribution of several storage compounds and intrinsic biomolecules in diverse eukaryotic and prokaryotic cells^74–77^.

In summary, we have developed an exceptionally high-throughput SRS-FISH platform and successfully applied this new tool to identify efficient mucosal sugar utilizers in the human gut microbiome. It is safe to assume that SRS-FISH can be applied to a broad range of environmental samples (e.g. marine sediments, soil) including those where some autofluorescence background is an issue because SRS is more resilient to sample autofluorescence than spontaneous Raman^78,79^. Meanwhile, SRS-FISH is not limited to microbiome samples. With the state of art SRS metabolism imaging and versatile FISH techniques, such as probing abnormal proliferation of chromosomes or targeting mRNA, SRS-FISH will be broadly applicable to eukaryotes. By allowing the scanning of multiple samples in a fast and sensitive manner, SRS-FISH is well-suited to reveal fine-scale temporal, individual, and spatial patterns in a variety of specimens, which can otherwise be missed by existing methods due to their low-throughput.

## Methods

### SRS-FISH platform

A dual output, 80-MHz femtosecond (fs) pulsed laser (InSight X3, Spectra-Physics, USA) provides the pump beam (tunable from 680 nm to 1300 nm) and the Stokes beam (fixed at 1045 nm) for the fs SRS system (**Fig. 1d**, left panel). Stimulated Raman loss (SRL) provides the SRS intensity by detecting the modulation transfer from the Stokes to the pump beam. The Stokes beam was modulated by an acousto-optic modulator (1205c, Isomet Corporation, USA) at ∼2.26 MHz. The two beams were then combined by the dichroic mirror and directed into a lab-built laser scanning microscope. A 60× water objective (UPlanApo 60XW, NA=1.2, Olympus, Japan) focused the collinear beams on the sample. The power on the sample was ∼6 mW for the pump beam and ∼32 mW for the Stokes beam. The 2D galvo scanning unit (6215H, Cambridge Technique, USA) was conjugated to the back aperture by a 4f-system and scanned the laser focus to create the SRS image. An oil condenser (NA=1.4, Aplanat Achromat 1.4, Olympus, Japan) alleviated cross-phase modulation induced background in SRS by fully collecting laser beams. The collected beams were filtered by two filters (HA825/150m, Chroma, USA) and only the pump beam was detected by the silicon photodiode (PD, S3994-01, Hamamatsu, Japan). Then the photon-converted electric signal was first separated into alternating current (AC) readout and direct current (DC) readout. Then the AC signal was amplified by the lab-build resonant amplifier circuit centered at ∼2.26 MHz. After that, the AC signal was further extracted by a lock-in amplifier (HF2LI, Zurich Instrument, Switzerland). The quantitative chemical maps were created when the energy difference between the pump and the Stokes beam matched the vibrational energy of the targeted chemical bond (C-D centered at 2168 cm^−1^ and C-H centered at 2946 cm^−1^) (**Fig. 1 b**). The off-resonance images were recorded when the pump beam was tuned to 830 nm (targeting 2479 cm^−1^) for subsequent background subtraction in SRS (**Supplementary Fig. 2**).

To incorporate FISH visualization into the platform, we implemented TPEF in the fs SRS system (**Fig. 1a**, right panel). Forward detection with a higher collection efficiency of the condenser better preserved the fluorescence signal. With a flip mirror, the light was directed into the fluorescence collection devices. Two SiPMs (C14455-3050GA, Hamamatsu, Japan) modules were implemented to provide better quality fluorescence images compared to photomultiplier tubes (H7422-40, Hamamatsu, Japan) with an external pre-amplifier^28^. A 75 mm focal length lens focused the emission light onto the SiPMs with a 605 nm cut-on dichroic mirror (DMLP605, Thorlabs, USA) that separated the emission into two paths. Two filters centered at 570 nm (ET570/20x, Chroma, USA) and 670 nm (ET670/50m, Chroma, USA) were used to detect the fluorescence from different FISH labeled cells with Cy3 or Cy5, which can be efficiently excited by the Stokes beam and the pump beam (for C-D, C-H or off-resonance due to the wide two photon absorption bandwidth) in SRS respectively. A data acquisition card (PCIe-6363, National Instruments, USA) collected the final output to construct the images.

To determine the SRS lateral resolution, 200 nm polymethyl methacrylate beads (MMA200, Degradex phosphorex, USA) were used. By deconvoluting the SRS image with simulated 200 nm size beads, we obtained the point spread function (PSF) of SRS^80^, which was the product of pump and Stokes beam and had a FWHM around 242.3 nm (**Supplementary Fig. 1**). With 210 nm yellow green fluorescence beads (17151-10, Polysciences, USA), the spatial and axial resolution of non-degenerate (ND)-TPEF, pump beam TPEF, and Stokes beam TPEF were measured in the liquid or dry conditions. The ND-TPEF was also the product of pump and Stokes beam, thus it is similar to SRS PSF measurement. Then ND-TPEF images were obtained by subtracting fluorescence images with pump and Stokes both on by pump only TPEF images and Stokes only TPEF images. Due to the broad two photon absorption, we could detect fluorescence using different laser beams, under both liquid and dry imaging conditions, although signal from the 1045 nm TPEF beam in dry conditions was too weak to see the fluorescence beads. The resolution was calculated following the steps of SRS resolution calculation. In the liquid environment, the lateral resolutions of pump only TPEF, Stokes only TPEF, and ND-TPEF by pump and Stokes were 300∼400 nm while all of them degraded 1.5∼2 times to 500 nm∼600 nm when imaging in the dry conditions. The axial resolutions degraded by 3 times from ∼1.8 µm in the liquid environment to ∼6 µm in the dry conditions (**Supplementary Fig. 1**). This measurement further demonstrated the degraded sensitivity of SRS when imaging in the dry conditions. Meanwhile, TPEF degradation in the dry conditions did not impede the precision of generating FISH masks (**Supplementary Fig. 1, Fig. 4c**), which also has a lower requirement of signal level.

To determine the limit of SRS detection, DMSO (≥99.9%, D8418-500ML, Sigma-Alorich) and DMSO-d6 (99.96 atom % D, 156914-1G, Sigma-Alorich) were used as the standard samples to calibrate the system. The slopes between SRS intensity and solution concentration were acquired by fitting concentrations between 0.1 and 4 M, which were 8.123 arb. Unit per M for DMSO-d6 and 14.099 arb. unit per M for DMSO. The standard deviation of pure water measured at C-D and C-H regions were 0.023 and 0.02. The limits of detection were acquired by three standard deviations divided by the slope multiply by six, due to six C-D or C-H bonds in one DMSO-d6 or DMSO molecule. Then the limits of SRS detection of C-D and C-H were 51 mM and 25.8 mM, corresponding to around five million C-D bonds and three million C-H bonds inside the SRS excitation volume.

### Image acquisition

Fixed cells were spotted onto a poly-L-lysine coated coverslip, and covered and sealed by another coverslip with a spacer in between^37^. Samples were prepared in this way to reduce cross-phase modulation signal while keeping the sample in the liquid environment that matches the refractive index of the water objective used for imaging. For imaging in the dry conditions, fixed cells were spotted onto a poly-L-lysine coated coverslip, dried, and then covered by another coverslip with spacers at two opposite sides. For each sample, three fields of view (FOV) were scanned by a motorized stage, or manually for SRS-FISH analysis. Three channels of SRS images (C-D, C-H, and off-resonance) and two fluorescence images (Cy3 and Cy5) were collected as a full image set for analyzing two populations targeted with FISH. Although fluorescence images could be acquired simultaneously with SRS-CD by splitting the output beam in the forward direction or collecting epi-fluorescence signal, limitations in the data acquisition card do not provide higher sampling speed for multichannel sampling. So all the images were acquired sequentially. Each FOV was 42.8^2^ or 85.6^2^ μm^2^ with 214 nm per step and covered around 300∼400 or 1200∼1600 cells. Depending on the signal intensity level, 10 μs pixel dwell time with 1∼10 frames average was applied for either SRS or fluorescence images. The laser wavelength tuning and stabilizing time for changing between different SRS frames was around 10 s. The throughput of SRS-FISH analysis is around 10-100 ms per cell (∼10-100 cells per second) by taking into account the FOV moving time and laser wavelength switching time. For the microbiome test of 3 individuals’ samples in 8 different conditions, 3 randomly selected field of views were measured, which totally covered around thirty thousand cells.

### Image processing

For bacterial cultures, three SRS images at C-D, C-H, and off-resonance were collected without fluorescence reference as their identities were known beforehand. For applying FISH masks in two targeted populations in the artificial mix or gut microbiome samples, two fluorescence images of Cy3 and Cy5 were recorded along with three SRS images. After all images were acquired, SRS images were first subtracted by the mean background intensity respectively to eliminate the signal from the glass substrate. Then three channels were rescaled according to the DC readout from the PD with resonant amplifier circuit (**Supplementary Fig. 3**). The AC and DC signals were linear to the pump power inside the power range used in this experiment. Thus, the subtraction between the rescaled C-D, C-H with off-resonance channels can eliminate the other pump-probe background.

Fluorescence images acquired in the dry conditions were aligned to SRS images acquired after water immersion to reduce the mask measurement error from sample drift caused by water immersion. C-H channels or fluorescence channels were used to generate the single cell measuring masks depending on the need and availability of the fluorescence images. The created masks enabled the measurement of single cell average level in off-resonance subtracted C-D and C-H images. The %CD_SRS_ of masked cells were then calculated as described in the Result. All statistical analyses were performed using MATLAB (The MathWorks, USA).

To circumvent the influence of food residues normally present in fecal samples on our results, the regions with overlapping signals in the Cy3 and Cy5 channels were removed from the measurement mask. Wide emission fluorescence is an indication of autofluorescence signal typically originated from food particles^81^. Regions of abnormally strong intensity in the SRS image can be also attributed to the photothermal signal or transient absorption of the easily burned food residue. Therefore, intensity outliers (>2 times of the average intensity) recorded for both C-D and C-H channels were also removed from the mask measurement results, such as the very bright spots in the negative control (H_2_O+Glc) C-D channel in **Fig. 5a**.

### Growth and labeling of microbial pure cultures

*Escherichia coli* K12 (DSM 498) was grown aerobically at 37°C in Luria-Bertani liquid medium (LB; DSMZ medium 381) or M9 minimal medium containing per L of medium: 7.5 g Na_2_HPO_4_.2H_2_O, 3 g KH_2_PO_4_, 0.5 g NaCl, 0.5 g NH_4_Cl, 1 mM MgSO_4_, 0.3 mM CaCl_2_, 1 μg thiamine hydrochloride (Sigma-Aldrich), 1 μg biotin (Sigma-Aldrich), trace elements, 0.4% (w/v) D-Glucose (Carl Roth GmbH). For D labeling of *E. coli* cultures, 50 μl of a stationary-phase culture were used to inoculate 5mL of LB or M9 medium containing different percentages (vol/vol) of D_2_O (99.9% atom % (at%) D; Sigma Aldrich). Cells were grown until the late exponential phase (3 h in LB medium or 8-10 h in M9 medium), harvested by centrifugation and fixed in 4% formaldehyde in phosphate-buffered saline (PBS) for 2 h at 4°C. Cells were subsequently washed once with PBS and stored at 4°C until further use. *Bacteroides thetaiotaomicron* (DSM 2079) cells were grown anaerobically (85% N_2_, 10% CO_2_, 5% H_2_) in BHI broth (DSMZ medium 215c) or in *Bacteroides* minimal medium^82^ containing different percentages (vol/vol) of D_2_O (99.9% atom % (at%) D; Sigma Aldrich). After 6 hours of growth at 37°C, cells were harvested by centrifugation, resuspended in PBS, and fixed by adding formaldehyde in PBS as described above. All cells were stored in a PBS solution at 4 °C until further use.

### Gut microbiome incubations

Human fecal samples were collected from three healthy adult individuals (one male and two females between the ages of 25 to 38) who had not received antibiotics in the prior 3 months. Sampling of human fecal samples was approved by the University of Vienna Ethics Committee (reference #00161). Study participants provided informed consent and self-sampled using an adhesive paper-based feces catcher (FecesCatcher, Tag Hemi, Zeijen, NL) and a sterile polypropylene tube with the attached sampling spoon (Sarstedt, Nümbrecht, DE). Samples were transferred into an anaerobic tent (Coy Laboratory Products, USA) within 30 min after sampling. Samples were suspended in M9 medium (prepared with H_2_O and without glucose) to achieve a concentration of 0.1 g.ml^−1^, and further diluted 20 times in this medium. The homogenate was left to settle for 10 minutes, and the supernatant was then distributed into glass vials. An equal volume of M9 (without glucose) prepared with either D_2_O (99.9% atom % (at%) D; Sigma Aldrich) or H_2_O (for the negative control) was added to each tube, and each vial was finally supplemented with different concentrations of mucosal sugar monosaccharides (N-acetylneuraminic acid: 2 mg.ml^−1^; N-acetylglucosamine: 5 mg.ml^−1^; fucose: 2.5 mg.ml^−1^; galactose: 5 mg.ml^−1^; N-acetylgalactosamine: 2.5 mg.ml^−1^), D-glucose (5 mg.ml^−1^) or nothing (no-amendment control) (all amendment chemicals were from Sigma–Aldrich, except D(+)-galactose which was purchased from Carl Roth GmbH). These concentrations were selected based on reported concentrations of the different monosaccharides in purified hog gastric mucin and mucin gels, as described in^17^. After incubation for 6 h at 37°C under anaerobic conditions (5% H_2_, 10% CO_2_, 85% N_2_), samples were pelleted, washed with 1× PBS to remove D_2_O and then fixed in 3% formaldehyde for 2 h at 4°C. Samples were finally washed two times with 1 ml of PBS and stored in PBS at 4°C until further use.

### Fluorescence *in situ* hybridization

Fixed cells (100 μl) were pelleted at 14000 *g* for 10 min, resuspended in 100 μl 96% analytical grade ethanol and incubated for 1 min at room temperature for dehydration. Subsequently, the samples were centrifuged at 14000*g* for 5 min, the ethanol was removed, and the cell pellet was air-dried. Cells were hybridized in solution (100 μl) for 3 h at 46°C. The hybridization buffer consisted of 900 mM NaCl, 20 mM TRIS HCl, 1 mM EDTA, 0.01% SDS and contained 100 ng of the respective fluorescently labeled oligonucleotide as well as the required formamide concentration to obtain stringent conditions (**Supplementary Table S1**). After hybridization, samples were immediately transferred into a centrifuge with a rotor pre-heated at 46°C and centrifuged at 14000 *g* for 15 min at maximum allowed temperature (40°C), to minimize unspecific probe binding. Samples were washed in a buffer of appropriate stringency^83^ for 15 min at 48°C, cells were centrifuged for 15 min at 14 000 g and finally resuspended in 20 μl of PBS. Cells (5 μl) were spotted on Poly-L-lysine coated cover glasses No. 1.5H (thickness of 170 μm ± 5 μm, Paul Marienfeld EN) and allowed to dry overnight at 4°C, protected from light. Excess of salt was removed by dipping the coverslips 2× in ice-cold Milli-Q water and allowed to dry at room temperature protected from light.

### Confocal fluorescence microscopy

Samples were spotted onto microscope slides (Paul Marienfeld EN) with Poly-L-lysine coating and visualized using an Olympus scanning confocal microscope (FV3000) with a 60x oil objective (PLAPON60XO, 1.42 NA, 0.15 mm WD, Olympus) and two high-sensitivity photomultiplier tubes (PMTs). Cy3 double-labeled probe Bac303 was excited by a 514 nm solid state diode laser and its emission was collected between 530 nm and 630 nm. Cy5 double-labeled probe Erec482 was excited by a 640 nm solid state diode laser and its emission was collected between 650 nm and 750 nm. Transmission images were also collected for validating the focus and the distribution of the whole gut microbiome sample. The ideal spatial resolution limit is around 221 nm∼275 nm. The scanning step size of the confocal image was set to 103.58 nm, which was smaller than half of the diffraction limit of both beams and ensured sampling spatial frequency higher than the Nyquist frequency for both excitation wavelengths. Each acquired field of view is 212^2^ µm^2^.

### Spontaneous Raman microspectroscopy

Fixed cells were spotted on aluminum-coated slides (Al136; EMF Corporation). Excess of salt was removed by dipping the slide twice into ice-cold Milli-Q water. Individual cells were observed under a 100×/0.75 NA microscope air objective Single microbial cell spectra were acquired using a LabRAM HR800 confocal Raman microscope (Horiba Jobin-Yvon) equipped with a 532-nm neodymium-yttrium aluminum garnet (Nd:YAG) laser and 300 grooves/mm diffraction grating. Spectra were acquired in the range of 400–3200 cm^−1^ for 30 s with 2.18 mW laser power. Spectra were then aligned according to the phenylalanine peak region and normalized by dividing the spectral intensity at each wavelength by the total spectral intensity. For quantification of the degree of D incorporation into the cells, %CD was calculated using the integration of the intensity of bands assigned to C–D (2040–2,300 cm^−1^) and C–H (2,800–3,100 cm^−1^)^10^. For Raman-FISH analyses, cells displaying signals originating from hybridized oligonucleotide probes were identified using epifluorescence imaging on the Raman confocal microscope equipped with a 100- W Xenon lamp, a standard Cy5 filter block and an F-View camera (Soft Imaging Systems). Raman spectra of targeted cells were obtained by switching image acquisition to the inbuilt Raman CCD detector.

## Data and codes availability

The data and codes that supports the findings of this study are available from the corresponding author upon request.

## Acknowledgements

Research reported in this publication was funded by National Institutes of Health under award Number R35GM136223 and R01AI141439 to J.-X.C. and supported by the Boston University Micro and Nano Imaging Facility and the Office of the Director, National Institutes of Health of the National Institutes of Health under award Number S10OD024993. The content is solely the responsibility of the authors and does not necessarily represent the official views of the National Institute of Health. Funding for the presented research was also provided by the Austrian Science Fund (FWF) via the Wittgensteinaward to M.W. (Z383-B) and via a Young Independent Research Group grant to F.C.P. (ZK-57). We thank Arno Schintlmeister for valuable comments on the manuscript.

## Author contributions

X.G., F.C.P., M.W. and J.-X.C. conceived and designed the study. X.G., F.C.P. and M.M. performed the experiments and analyzed the data. X.G., F.C.P., M.W. and J.-X.C. wrote the manuscript. D.B. provided support and input to the microbiome experiments. M. Z. provided input on D labeling and cell fixation protocols. All authors have given approval to the final version of the paper.

## Competing Interests

The authors declare no competing interests.

## Notes

### Competing Interest Statement

The authors have declared no competing interest.

