## Supplementary for "SRS-FISH: High-Throughput Platform Linking Microbiome Function to Identity at the Single Cell Level"

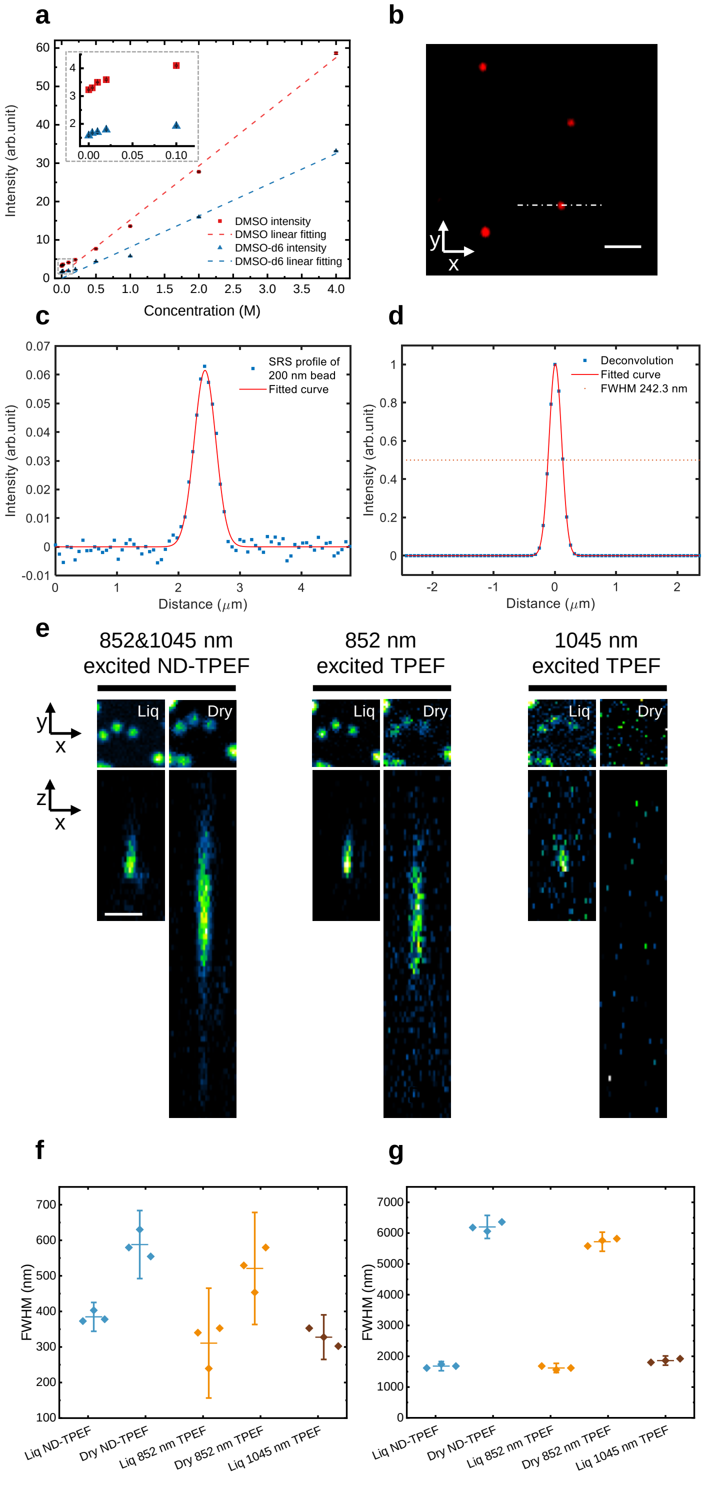


**Supplementary Figure 1.** Femtosecond SRS system characterization of sensitivity and resolution. **a**, Femtosecond SRS system sensitivity test with increasing concentrations of dimethyl sulfoxide (DMSO) and deuterated DMSO (DMSO-d6). The inserted grey dashed box at left top corresponds to the magnification of content in the grey dashed box at the left bottom. The lowest detectable concentration is 25.8 mM C-H bonds according to 4.3 mM DMSO and 51 mM C-D bonds according to 8.5 mM DMSO-d6. **b**, SRS imaging of 200 nm polymethyl methacrylate beads at C-H channel in lateral direction. Scale bar: 2 μm. **c**, Cross section profile of the dash dot line across the bead in **b**. **d**, lateral point spread function (PSF) by deconvoluting the SRS image with simulated 200 nm beads. **e**, TPEF imaging of 210 nm yellow green fluorescence beads with different excitations under the liquid (Liq) or dry conditions in lateral or axial directions. ND: non-degenerate. Scale bar: 2 μm. **f**, lateral and **g**, axial PSF full width at half maximum (FWHM) under different excitation and sample conditions. Interval plots in **f**, **g** represent the mean, with the extended lines represent the value within 95% confidence interval. Diamond markers represent individual measurements.

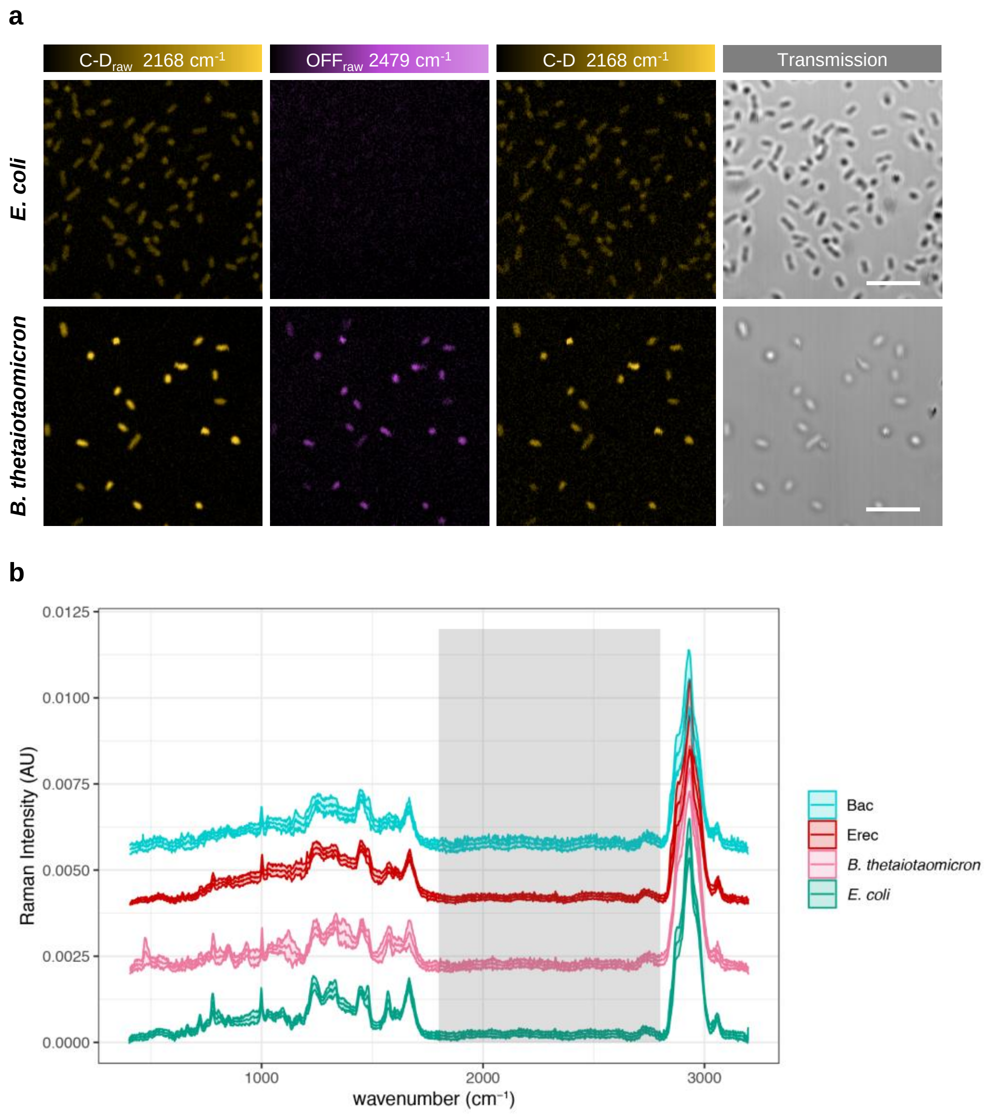


**Supplementary Figure 2. a**, SRS imaging of C-D (yellow, raw and after off-resonance subtraction) and off-resonance (purple) frequencies of hybridized *E. coli* (grown in 50% $D_2O$-containing M9 medium) or *B. thetaiotaomicron* (grown in 50% $D_2O$-containing *Bacteroides* minimal medium) cells when imaged in liquid environment. Scale bar, 10 µm. **b**, Spontaneous Raman spectra of hybridized *E. coli* (green) and *B. thetaiotaomicron* (pink) cells from pure cultures grown in $H_2O$-containing M9 or $H_2O$-containing *Bacteroides* minimal medium, respectively. In the same chart, spectra from cells targeted by Erec482-Cy5 (Erec, in red) and Bac303-Cy5 (Bac, in cyan) oligonucleotide probes within a gut microbiome sample incubated in $H_2O$-containing M9 medium are also shown. Lines depict mean values and shaded regions depict standard deviations of spectra obtained from 9-21 measured cells. The silent region between 1800 and 2800 $cm^{-1}$ is highlighted in grey.

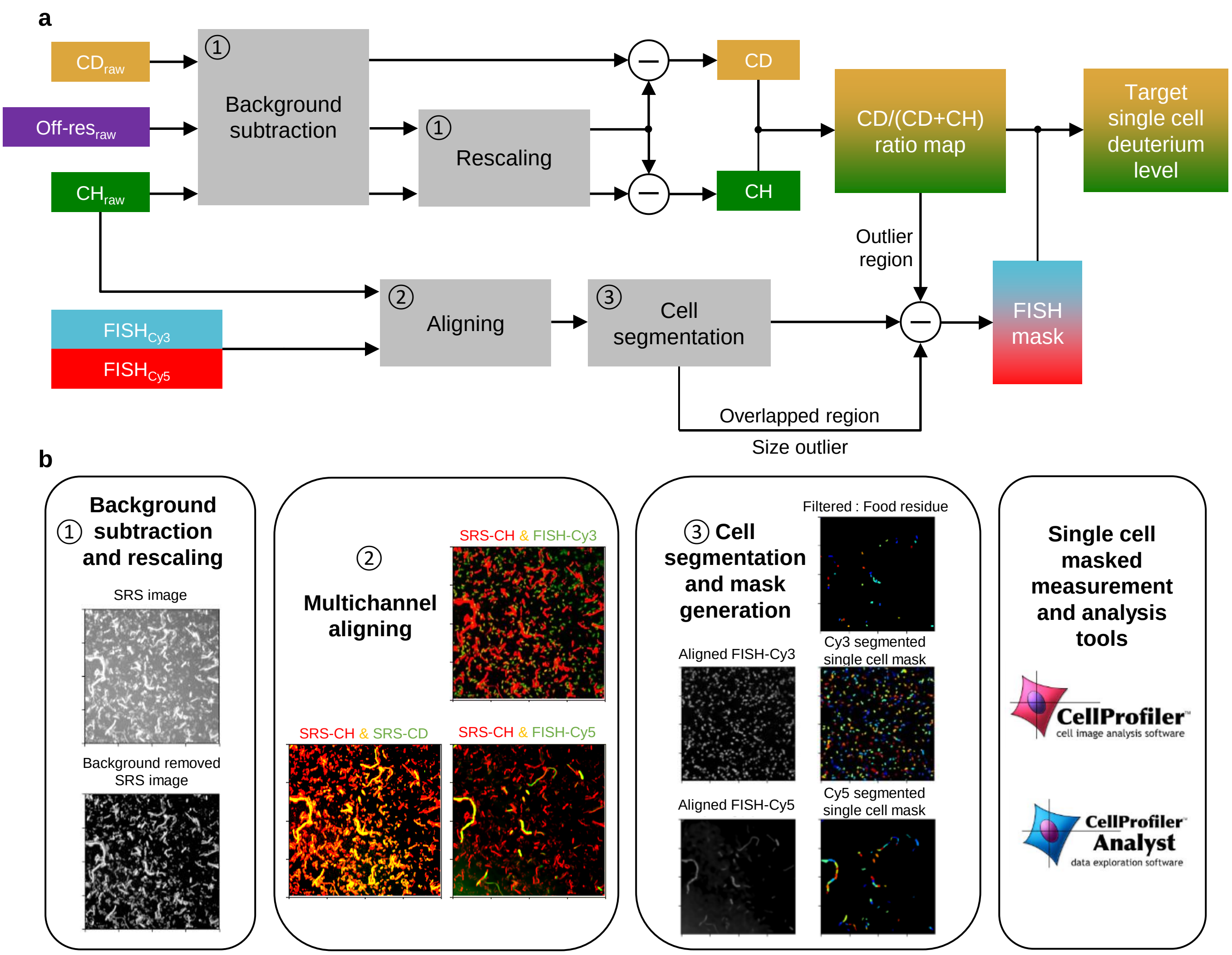


**Supplementary Figure 3.** Schematic representation of the strategy followed for calculation of single-cell cellular deuterium levels. **a**, Image processing flowchart. Minus sign stands for subtraction operation. **b**, Operations on gut microbiome of grey boxes in **a**. Detailed explanation is presented in Methods.

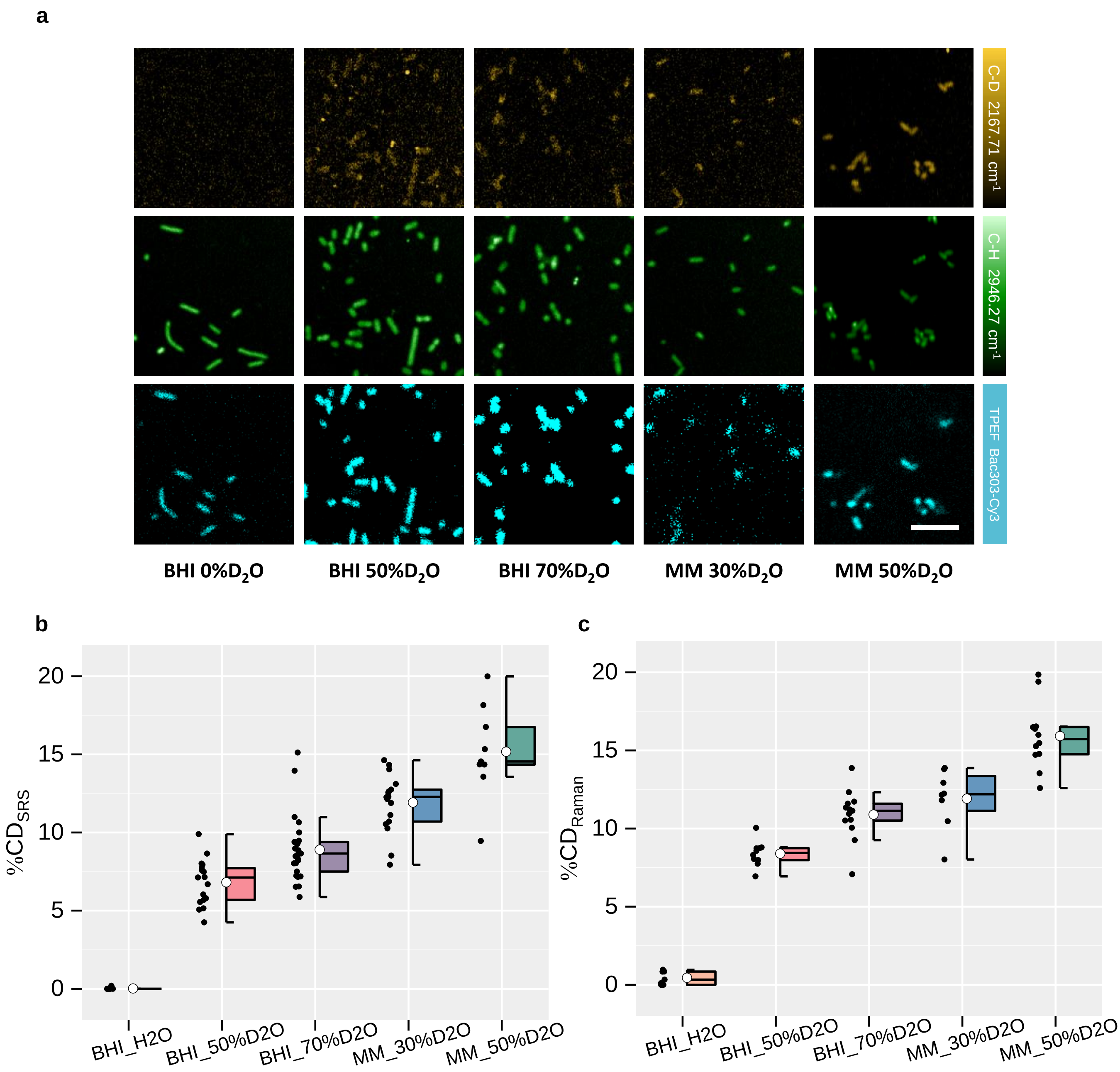


**Supplementary Figure 4.** SRS-FISH platform detects $D_2O$ incorporation by metabolically-active *B. thetaiotaomicron* cells. **a**, SRS imaging of C-D (yellow) and C-H (green) intensities of single *B. thetaiotaomicron* cells grown in BHI or in *Bacteroides* minimal media (MM) containing different percentages of $D_2O$, as indicated. FISH signal from *B. thetaiotaomicron* cells hybridized with a Bac303-Cy3 probe and acquired using the TPEF system is shown in cyan. Scale bar: 7 µm. Image contrast: C-D channel, min 0.2 max 2.5 (arb. unit); C-H channel, min 0 max 10 (arb. unit). Pixel dwell time: 50 µs. For details regarding data processing please refer to Supplementary Figure 3. Single cell C-D level distribution in *B. thetaiotaomicron* cells grown under the conditions described in **a**, measured with either SRS (**b**) or with spontaneous Raman microspectroscopy (**c**). In **b**-**c**, each dot represents a cell. Boxes represent median, first and third quartile. Whiskers extend to the highest and lowest values that are within one and a half times the interquartile range. The white circle in the middle of the box represents the mean value of the data.

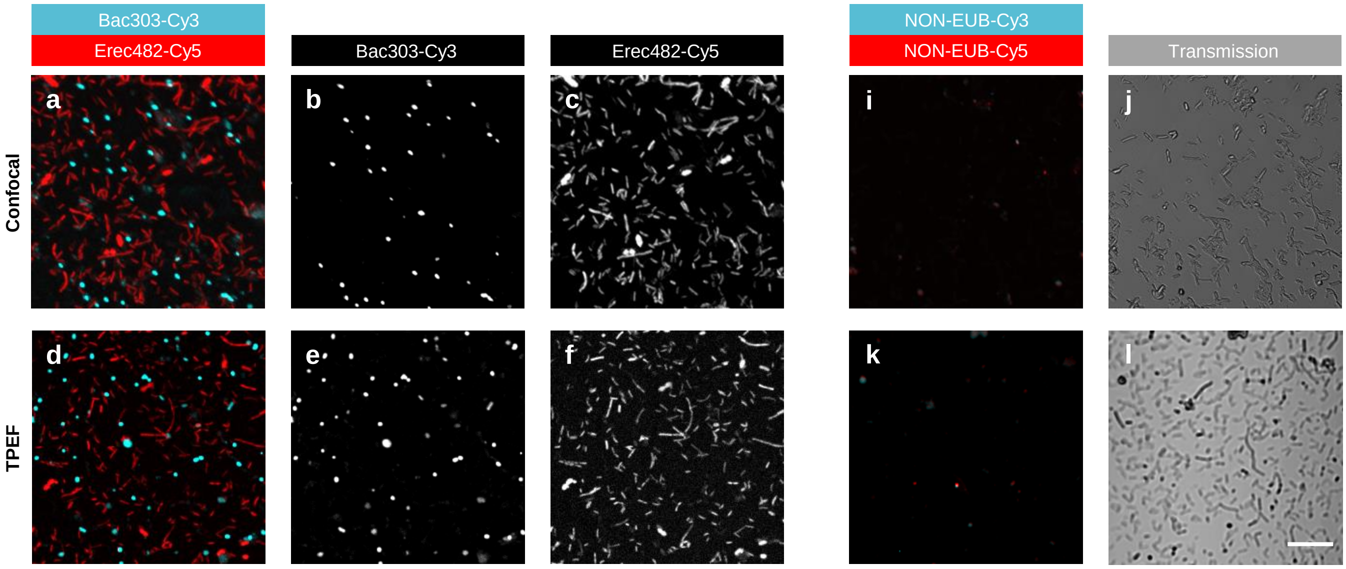


g

| FOV | Bac303-Cy3: Bacteroidales | Erec482-Cy5: Clostridia |
|---|---|---|
| 1 | 622 (21.9%) | 2224 (78.1%) |
| 2 | 538 (19.7%) | 2194 (80.3%) |
| 3 | 702(22.4%) | 2420 (77.6%) |
| Total % | 22.9% | 77.1% |

h

| FOV | Bac303-Cy3: Bacteroidales | Erec482-Cy5: Clostridia |
|---|---|---|
| 1 | 63 (16.4%) | 322 (83.6%) |
| 2 | 54 (14.7%) | 314 (85.3%) |
| 3 | 133 (22.8%) | 450 (77.2%) |
| Total % | 18.7% | 81.3% |

**Supplementary Figure 5.** Imaging of hybridized, fluorescently-labeled human gut microbiome samples by confocal and TPEF scanning microscopy. **a**-**g,** Representative images of the same microbiome sample hybridized with the probes Bac303-Cy3 and Erec482-Cy5 and imaged by confocal microscopy (**a**-**c)** or by TPEF (**d-f**)**. a**, **d**, Overlapped FISH signal under confocal (**a**) or TPEF (**d**) microscopy of Bac303-Cy3 (**b**, **e**) and Erec482-Cy5 (**c**, **f**) labeled cells. **g**, **h**, Percentage of cells from each population (cyan-labeled or red-labeled cells divided by the number of total cyan+red cells) for 3 different fields of view (FOV), under confocal (**g**) or TPEF (**h**) microscopy. **i**-**l,** Representative images of microbiome samples hybridized with the probes NON-EUB-Cy3 and NON-EUB-Cy5 (negative control probes employed to rule out non-specific binding of the oligonucleotide probes to cells) and imaged by confocal microscopy (**i, j)** or by TPEF (**k, l**). **i**, **k**, Overlapped fluorescence signal under confocal (**i**) or TPEF (**k**) microscopy of Cy3 and Cy5 channels. **j**, **l**, transmission images of **i** and **k** respectively. Scale bar: 15 µm.

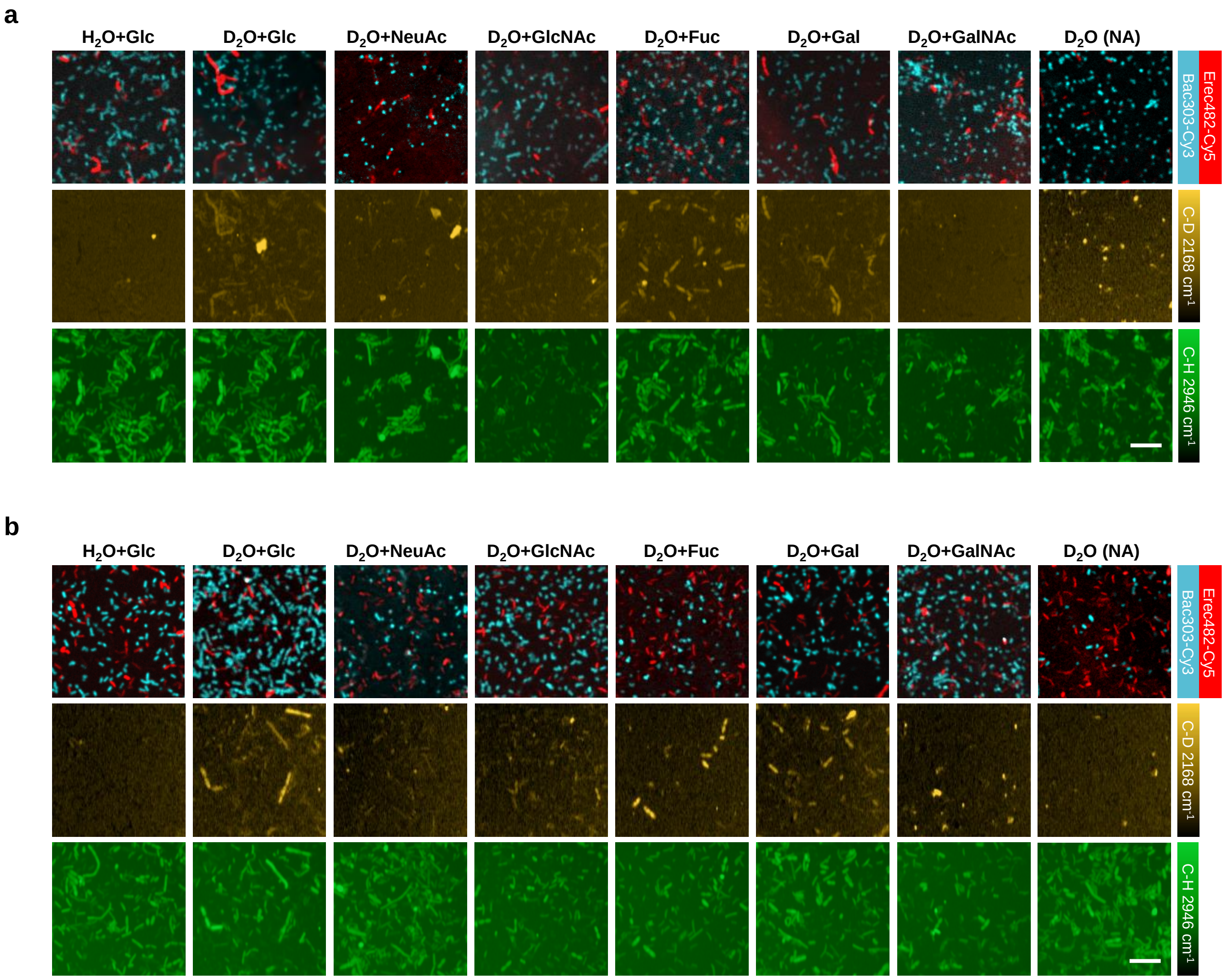


**Supplementary Figure 6.** Microbiome samples of volunteers 2 (**a**) and 3 (**b**) were incubated with different mucosal sugars and hybridized with the oligonucleotide probes Bac303-Cy3 and Erec482-Cy5. Representative images obtained by TPEF (top row) and SRS (C-D middle row, C-H bottom row) are shown. Negative control: $H_2O$+Glucose. Positive control: $D_2O$+Glucose. NA: no amendment. Scale bar, 10 µm. For details regarding data processing please refer to Supplementary Figure 3.

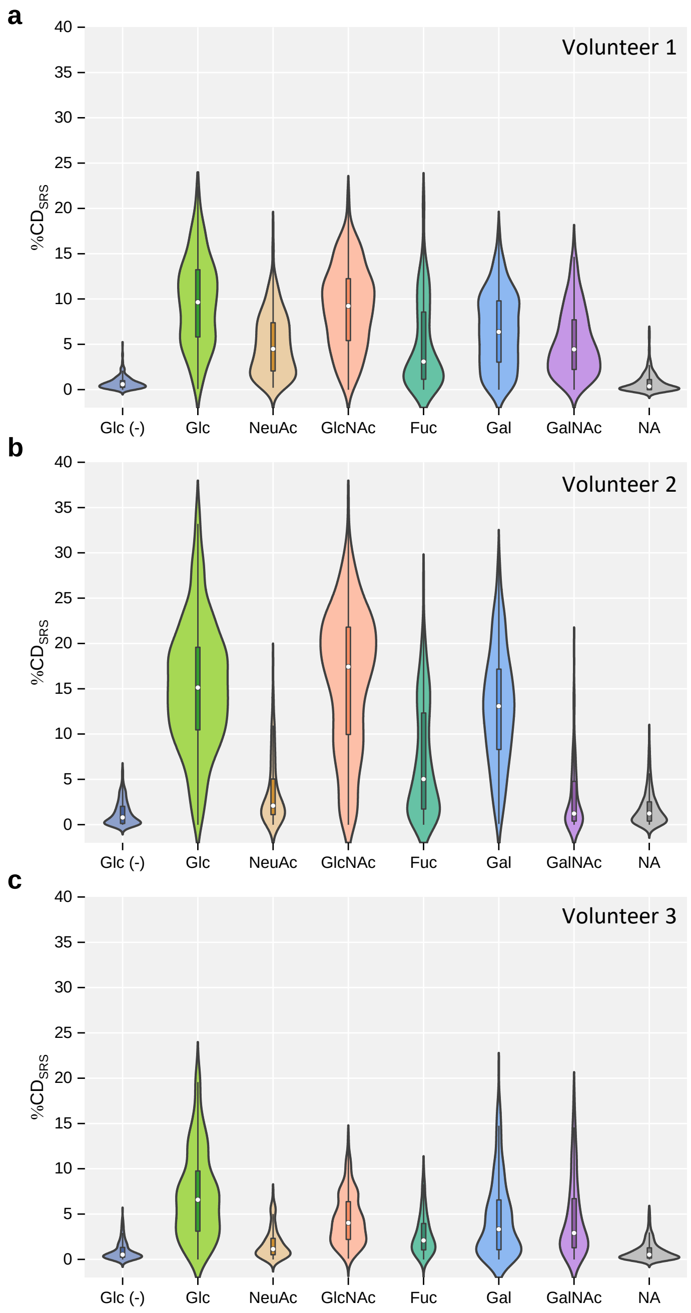


**Supplementary Figure 7.** Single cell C-D level distribution of all bacteria in the randomly selected field of views and amended with different mucosal sugars (and respective controls), for the same three volunteers shown in Figure 5. The measurement masks for the SRS C-D levels were created with SRS C-H channel. The violin shapes represent the distribution of cells. Boxes represent first and third quartile. White dots represent median value. Whiskers extend to the highest and lowest values that are within one and a half times the interquartile range.

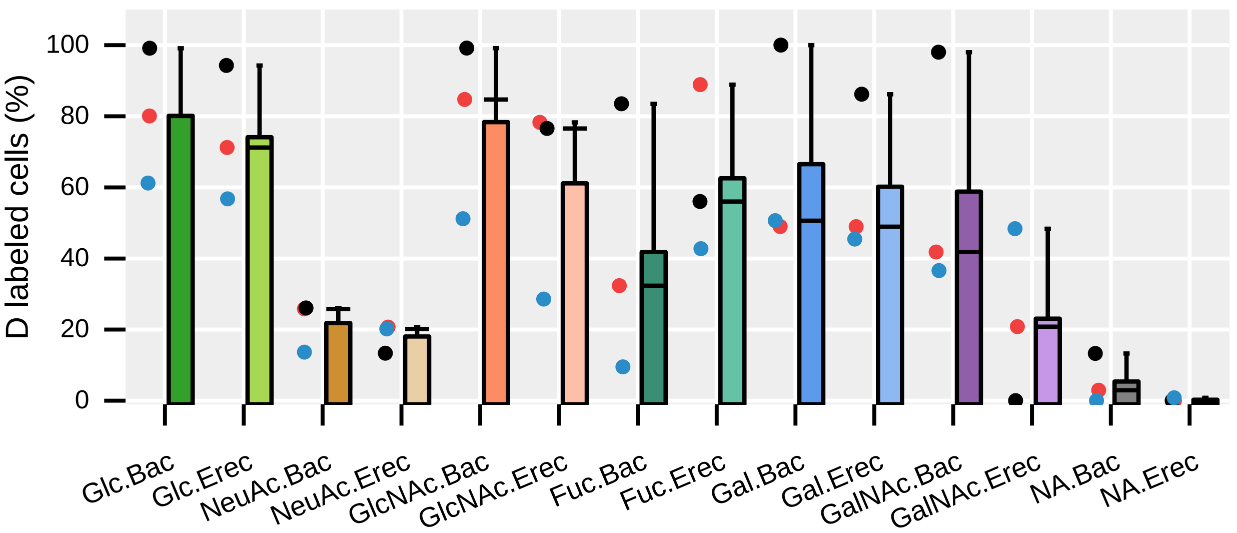


**Supplementary Figure 8.** Percentage of cells that show D incorporation above the threshold, determined as described in Figure 5. Data for the two abundant gut microbiome taxa (Bac and Erec) is shown for each amended sugar (and no amendment control). Each dot represents the percentage obtained for each of the three volunteers. Red dots: volunteer 1. Black dots: volunteer 2. Blue dots: volunteer 3. Bars represent mean. The intermediate lines represent the median. Whiskers extend to the values that are within one and a half times the interquartile range. Bac: Bac303-Cy3 targeted cells; Erec: Erec482-Cy5 targeted cells.

**Supplementary Table 1.** FISH probes for single cell analyses.

| Probe Name | Sequence (5' to 3') | Formamide (%) | Major target taxa[a] | Reference |
|---|---|---|---|---|
| **Gam42a** | GCCTTCCCACTTCGTTT | 35 | Class Gammaproteobacteria (28%) | [1] |
| **Erec482** | GCTTCTTAGTCARGTACCG | 10 | Order Lachnospirales (74%)<br>Family Lachnospiraceae (75%) | [2] |
| **Bac303** | CCAATGTGGGGGACCTT | 10 | Order Bacteroidales (39%)<br>Family Bacteroidaceae (86%)<br>Family Prevotellaceae (73%)<br>Family Barnesiellaceae (57%)<br>Family Tannerellaceae (50%) | [3] |
| **NON-EUB** | ACTCCTACGGGAGGCAGC | - | negative control probe complementary to EUB338 | [4] |

[a] According to Silva database v138.1 (https://www.arb-silva.de/). The probe coverage for each taxon is shown in parenthesis. Only taxa commonly present in the human gut are presented.


[1] Manz, W., Amman, R., Ludwig, W., Wagner, M. & Schleifer, K.-H. Phylogenetic Oligodeoxynucleotide Probes for the Major Subclasses of Proteobacteria: Problems and Solutions. *Syst. Appl. Microbiol.* **15**, 593–600 (1992).

[2] Franks, A. H. *et al.* Variations of Bacterial Populations in Human Feces Measured by Fluorescent In Situ Hybridization with Group-Specific 16S rRNA-Targeted Oligonucleotide Probes. *Appl. Environ. Microbiol.* **64**, 3336–3345 (1998).

[3] Manz, W., Amann, R., Ludwig, W., Vancanneyt, M. & Schleifer, K. H. Application of a suite of 16S rRNA-specific oligonucleotide probes designed to investigate bacteria of the phylum cytophaga-flavobacter-bacteroides in the natural environment. *Microbiol. Read. Engl.* **142** 1097–1106 (1996).

[4] Wallner, G., Amann, R. & Beisker, W. Optimizing fluorescent in situ hybridization with rRNA-targeted oligonucleotide probes for flow cytometric identification of microorganisms. *Cytometry* **14**, 136–143 (1993).

**Supplementary Table 2.** Statistical analysis results (expressed as P-value, two-sided Mann-Whitney U test) obtained when comparing CD levels for each sugar amendment condition with the CD levels of the negative control (Glc+$H_2O$). Bac: Bac303-Cy3 labeled cells. Erec: Erec482-Cy5 labeled cells. Grey background refers to $p>0.05$ and thus where CD levels do not significantly differ from the negative control group (Glc $H_2O$).

| Sugar amendment | Volunteer 1 | | Volunteer 2 | | Volunteer 3 | |
|---|---|---|---|---|---|---|
| | Bac | Erec | Bac | Erec | Bac | Erec |
| **Glc (+)** | $3.51\times10^{-40}$ | $1.03\times10^{-8}$ | $6.81\times10^{-86}$ | $2.54\times10^{-16}$ | $1.88\times10^{-12}$ | $2.18\times10^{-7}$ |
| **NeuAc** | $3.62\times10^{-2}$ | $1.00\times10^{-3}$ | $3.54\times10^{-11}$ | $1.20\times10^{-3}$ | $1.38\times10^{-2}$ | $3.81\times10^{-2}$ |
| **GlcNAc** | $5.18\times10^{-26}$ | $8.86\times10^{-8}$ | $6.57\times10^{-75}$ | $2.89\times10^{-13}$ | $1.38\times10^{-20}$ | $5.80\times10^{-3}$ |
| **Fuc** | $5.85\times10^{-6}$ | $4.14\times10^{-9}$ | $1.26\times10^{-40}$ | $1.75\times10^{-11}$ | $1.60\times10^{-3}$ | $4.27\times10^{-8}$ |
| **Gal** | $8.39\times10^{-17}$ | $7.73\times10^{-4}$ | $2.08\times10^{-61}$ | $1.33\times10^{-11}$ | $2.39\times10^{-14}$ | $2.34\times10^{-6}$ |
| **GalNAc** | $2.26\times10^{-8}$ | 0.254 | $1.09\times10^{-54}$ | $2.81\times10^{-2}$ | $2.89\times10^{-12}$ | $2.45\times10^{-6}$ |
| **NA** | 0.860 | 0.949 | 0.360 | 0.752 | 0.162 | 0.404 |
